# Post-Critical Period Transcriptional and Physiological Adaptations of Thalamocortical Connections after Sensory Loss

**DOI:** 10.1101/2024.11.19.624130

**Authors:** Laxmi Iyer, Kory Johnson, Sean Collier, Alan P. Koretsky, Emily Petrus

## Abstract

Graphical Abstract

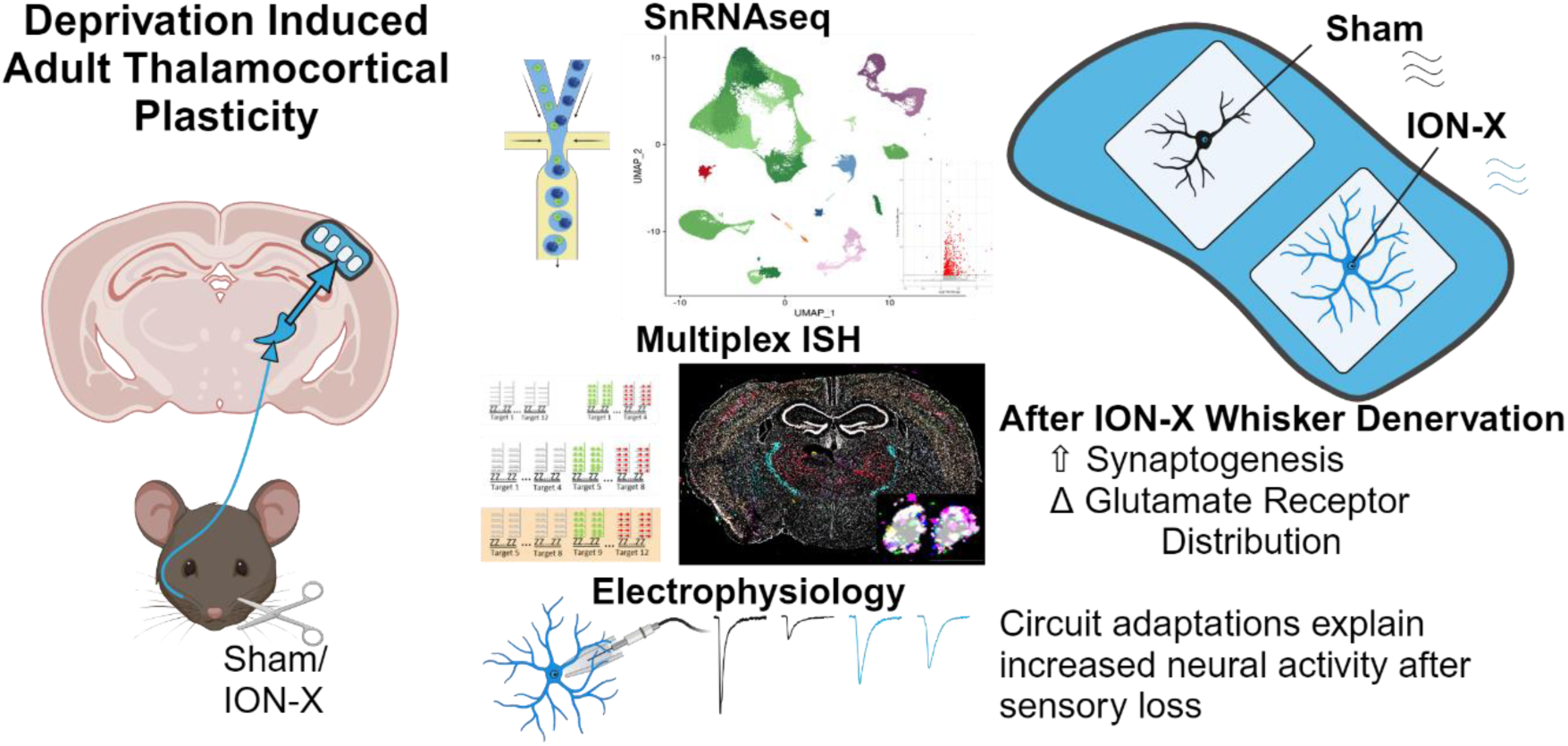

Unilateral whisker denervation activates plasticity mechanisms and circuit adaptations in adults. Single nucleus RNA sequencing and multiplex fluorescence in situ hybridization revealed differentially expressed genes related to altered glutamate receptor distributions and synaptogenesis in thalamocortical (TC) recipient layer 4 (L4) neurons of the sensory cortex, specifically those receiving input from the intact whiskers after whisker denervation.

Electrophysiology detected increased spontaneous excitatory events at L4 neurons, confirming an increase in synaptic connections. Elevated expression levels of Gria2 mRNA and functional GluA2 subunit of AMPA receptors at the TC synapse indicate the presence of stabilized and potentiated TC synapses to L4 excitatory neurons along the intact pathway after unilateral whisker denervation. These adaptations likely underlie the increased cortical activity observed in rodents during intact whisker sensation after unilateral whisker denervation. Our findings provide new insights into the mechanisms by which the adult brain supports recovery after unilateral sensory loss.

## Introduction

The adult brain retains the capacity to remodel after changes in sensory experience. Injuries to sensory systems cause a loss in sensory/motor function and lead to widespread cortical reorganization throughout sensory/motor circuitry^1–4^. Adaptive plasticity may support functional recovery by enhancing activity in intact brain regions involved in sensory processing and motor function^5–7^. Conversely, maladaptive plasticity can impair recovery, triggering pain phenotypes like hyperalgesia or impaired motor function^3,8–10^. It is unknown how these adaptations occur, thus predicting how individuals respond to unilateral sensory loss remains impossible. In addition, treatments to enhance recovery or reduce pain phenotypes remain ineffective due to a poor understanding of the mechanisms by which these circuit changes occur. We hypothesized that transcriptional profiling would identify the mechanisms by which sensory systems adapt after unilateral sensory loss. To study this response, a unilateral whisker denervation (infraorbital nerve transection: ION-X) was performed in mice. Previous findings identified a potentiated response in the intact primary somatosensory barrel cortex (S1BC) during stimulation of intact whiskers^11,12^, with dramatic plasticity occurring in the thalamocortical (TC) connections ^13,14^. This whisker denervation model allows for circuit and molecular interrogation of how the brain changes after unilateral sensory loss.

Experience-driven plasticity shapes circuits and their functions in the adult brain^13,14^. Historically, some components of the whisker sensory circuit were thought to be plastic only during the first post-natal week “critical period”, in particular at thalamocortical (TC) connections to layer 4 (L4) neurons^15,16^. However, recent studies have shown that changes in experience, driven by injury or sensory deprivation, can revive plasticity capacity in L4 neurons receiving TC input in multiple sensory systems^11,12,17–19^. This reopened window for plasticity in adults provides an opportunity to harness these mechanisms, potentially supporting recovery while animals and humans adapt to changes in sensory experience. In this study, we employed single nucleus RNA sequencing (snRNA-seq) to determine the mechanisms underlying the reopening of TC plasticity and the potentiation of cortical activity after ION-X.

It is unknown how the loss of whisker inputs via ION-X would alter gene expression and downstream protein functions within sensory circuitry. We used transcriptional profiling of L4 neurons to detect functional changes that occur in the adult mouse brain after unilateral sensory loss, and reveal additional mechanisms that boost plasticity. SnRNA-seq identified transcriptional adaptations in L4 neurons, which were validated with multiplex fluorescent in situ hybridization (mFISH) and whole-cell electrophysiological recordings.

Our integrative approach, combining molecular profiling with functional assays, uncovered a compelling link between gene expression changes, increased synaptic activity, and function after ION-X. Specifically, the identification of differentially expressed genes (DEGs) and pathways involved in glutamate receptor activity and synaptogenesis highlights the mechanism by which the increased neural activity is observed in people after unilateral injury resulting in sensory loss^12,20,21^. This approach provides a multi-level approach to validate snRNA-seq results with mFISH and electrophysiology. This study has identified the mechanisms that support cortical plasticity after injury-induced sensory loss; these findings may support the development of targeted interventions to improve recovery outcomes for individuals after unilateral denervation injuries.

## Results

### Transcriptional Profile of Intact Mouse Sensory Cortex

Intact S1BC tissue was collected 12 days after sham or ION-X surgery in mice (**Figure 1A**). This timepoint was chosen because gene expression is predicted to precede the maximal potentiation of the intact S1BC’s response to injury^11,19^. SnRNA-seq was performed, and the data were filtered to remove nuclei with low-quality reads, and >5% mitochondrial genes.

**Figure 1.**
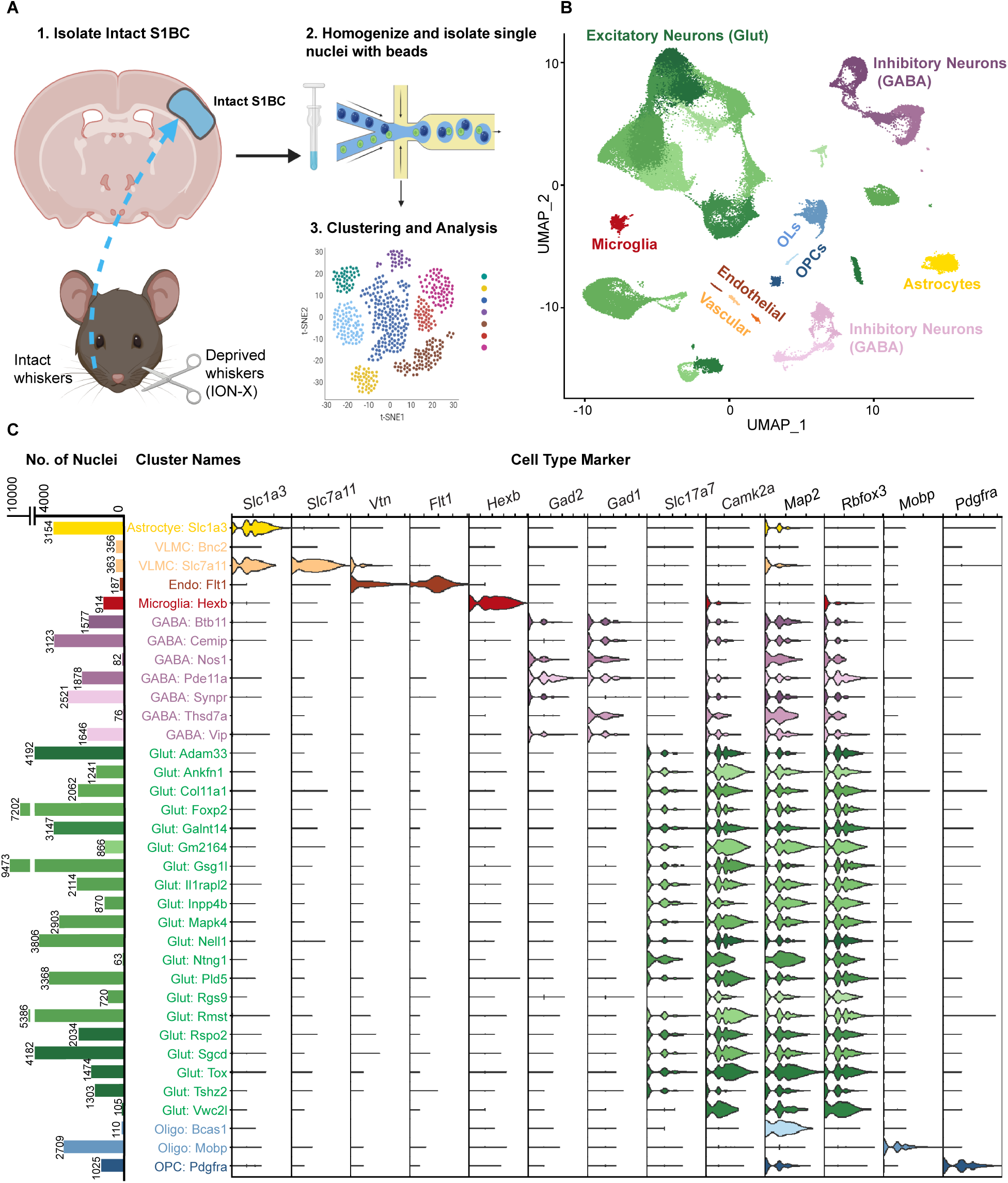
Transcriptional Profile of Intact Mouse Sensory Cortex. A) Schematic of SnRNA-seq workflow (n=8 C57BL6/J mice/group). B) Uniform Manifold Approximation and Projection (UMAP) with 83,390 nuclei clustered into 35 clusters, colored and annotated by cell type. C) Left: numbers of nuclei are shown for each cluster. Right: cell type-based classification was based on gene expression. See also Figures S1, S2 and Table S1.

Additionally, doublets were removed to ensure high-quality nuclei were used for further analysis (**Figure S1A-C, Methods**). After filtering, a total of 83,390 nuclei remained for downstream analysis and were grouped into 35 distinct clusters based on their gene expression profiles (**Figure 1C, Figure S2A**). The nuclei from these clusters were categorized into major cell types according to their expression of canonical cell type marker genes: astrocytes (*Slc1a3*), excitatory neurons (*Slc17a7, Camk2a, Rbfox3, Map2*), inhibitory neurons (*Gad1, Gad2*, *Rbfox3, Map2*), microglia (*Hexb*), endothelial and vascular cells (*Flt1, Vtn, Slc7a11*), oligodendrocytes (*Mobp*), and oligodendrocyte progenitor cells (*Pdgfra*) (**Figure 1B, C**). The distribution of nuclei across these clusters remained consistent between sham and ION-X samples, indicating no significant shifts occurred in the major cell types populations following injury (**Figure S1D-F, Table S1**). To further investigate the role of excitatory neurons and their potential involvement in post-injury cortical plasticity, we isolated clusters expressing canonical excitatory neuron markers (*Rbfox3, Map2, Slc17a7*), and re-clustered these nuclei (**Figure S2B**).

### Gene Expression Excitatory Layer 4 Neurons Detect Multiple Plasticity Pathways

A total of 76,232 nuclei from excitatory neurons were classified into cortical layers using auto-annotation of cortical layer marker genes. 26 clusters were automatically annotated with neurons assigned to different cortical layers; however, none of these clusters were specifically assigned to L4 (**Figure S3A, Table S2**). To improve classification accuracy, we manually curated a list of cortical layer marker genes derived from the Allen Brain Institute(24), annotated these 26 clusters, and assigned them to six cortical layers (**Figure 2A, S3B**, **Table S3**). To identify excitatory neurons in L4, we targeted clusters that expressed the pan-neuronal marker *Map2*, the excitatory cell marker *Slc17a7* (encodes for VGlut1), and the L4-specific marker *Rorb*^23^, (**Figure 2B**). This combination of markers enabled the identification of L4 excitatory neurons in the snRNA-seq data set. Multiplex fluorescent in situ hybridization (mFISH) was performed to detect *Map2, Slc17a7, and Rorb* in S1BC tissue, confirming that these Rorb-positive nuclei are located in L4 (**Figure 2C**). These *Map2/Slc17a7/Rorb* positive clusters were subsetted out of the snRNA-seq data set and re-clustered to identify three distinct subgroups of L4 excitatory neurons with unique, nonoverlapping markers of gene expression (**Figure S4A-B, Table S4-S5**). mFISH was performed on intact S1BC tissue but the specificity of these markers was less robust, with up to 25% of nuclei having overlapping gene expression (**Figure S4C-E**). In subsequent experiments all *Map2/Slc17a7/Rorb* positive nuclei were considered L4 excitatory neurons. Next, we examined differentially expressed genes (DEGs) in L4 excitatory neurons between sham and ION-X. 403 genes were significantly upregulated (adjusted p < 0.05) in L4 excitatory neurons in ION-X compared to sham mice (**Figure 2D, Table S6**). Gene Ontology enrichment analysis detected upregulated pathways associated with synapse assembly and organization, cell-cell adhesion, and glutamate receptor signaling (**Figure 2E, S4F-G**). Genes associated with plasticity and circuit remodeling are upregulated in L4 excitatory neurons, which receive input from the intact whisker set after unilateral sensory loss. These neurons likely direct circuit plasticity to support circuit remodeling after unilateral sensory loss.

**Figure 2.**
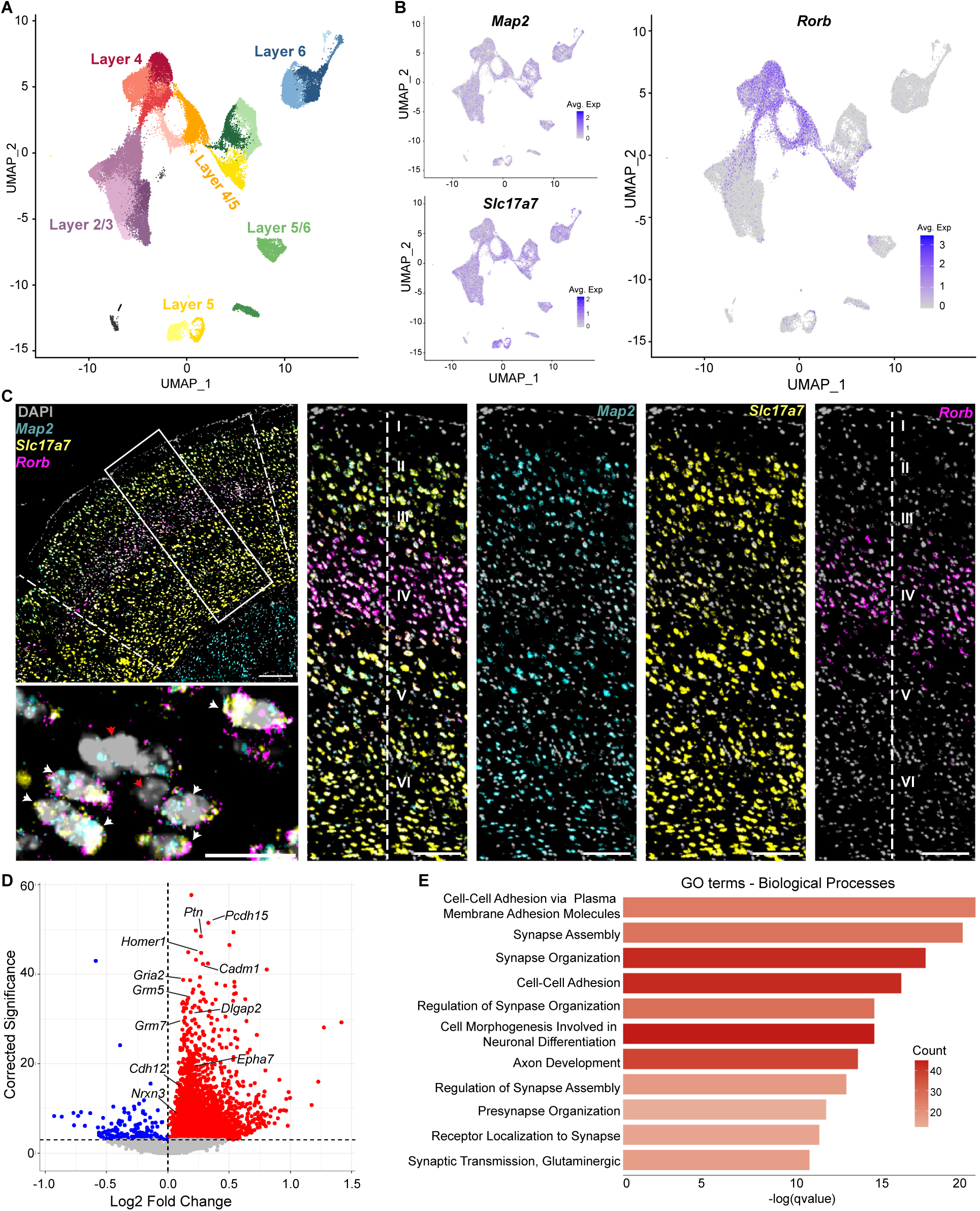
Transcriptional Characterization of Excitatory Layer 4 (L4) Neurons. A) UMAP of 76,232 excitatory neuronal nuclei colored and annotated by cortical layers. B) Feature plots of neuronal marker *Map2*, excitatory cell maker *Slc17a7,* and L4 marker *Rorb* were used to identify L4 excitatory neurons. C) mFISH detects neuronal marker gene *Map2*, excitatory cell marker *Slc17a7*, and L4 marker *Rorb*; counter-stained with DAPI to identify L4 excitatory neurons in sensory cortex. Representative higher magnification images show cortical layers in a composite and separate image for each gene. The bottom left image shows individual cells identified as L4 excitatory neurons with white arrows positive for all 3 markers and red arrows negative. Scale bars: 200µm on the left top; 10 µm left bottom; 100 µm for all right. D) Volcano plot showing differentially expressed genes (DEGs) in L4 neurons. E) Gene Ontology enrichment analysis identified upregulated biological processes ranked by q-value and colored according to the number of DEGs. See also Figures S3, S4 and Table S2,S3,S6.

*Altered Glutamate Receptor Signaling in the Intact Sensory Pathway after ION-X* Upregulated genes and pathways in L4 excitatory neurons were related to glutamate receptor function and synapse formation. First, we focused on 3 genes related to glutamate receptor function that were upregulated after ION-X in L4 excitatory neurons: *Homer1, Grm5,* and *Gria2* (adjusted p = 1.18×10^-15^, 2.76×10^-11^, 4.97×10^-13^ respectively). *Homer1* is a scaffolding protein that regulates metabotropic glutamate receptor (mGluR) anchoring and function, including mGluR5 (encoded by *Grm5* mRNA). *Gria2* encodes for the GluA2 subunit of ionotropic glutamate AMPA receptors (AMPARs) which regulates calcium entry into neurons during synaptic activity. We quantified their expression patterns using mFISH in nuclei that were triple positive for *Map2/Slc17a7/Rorb* (**Figure 3A, B**). Gene expression quantification was measured as both the percentage of positive nuclei and the number of mRNA puncta per nucleus. Consistent with snRNA-seq results, genes that were upregulated after ION-X were also increased when measured by mFISH (**Figure 3C, Table S7**, *Homer1*; % positive nuclei: t-test p=0.0011; *Grm5*; % positive nuclei: t-test p=0.001; average puncta p=0.0158; *Gria2*; % positive nuclei: t-test p=0.0013; average puncta p=0.0216;).

**Figure 3.**
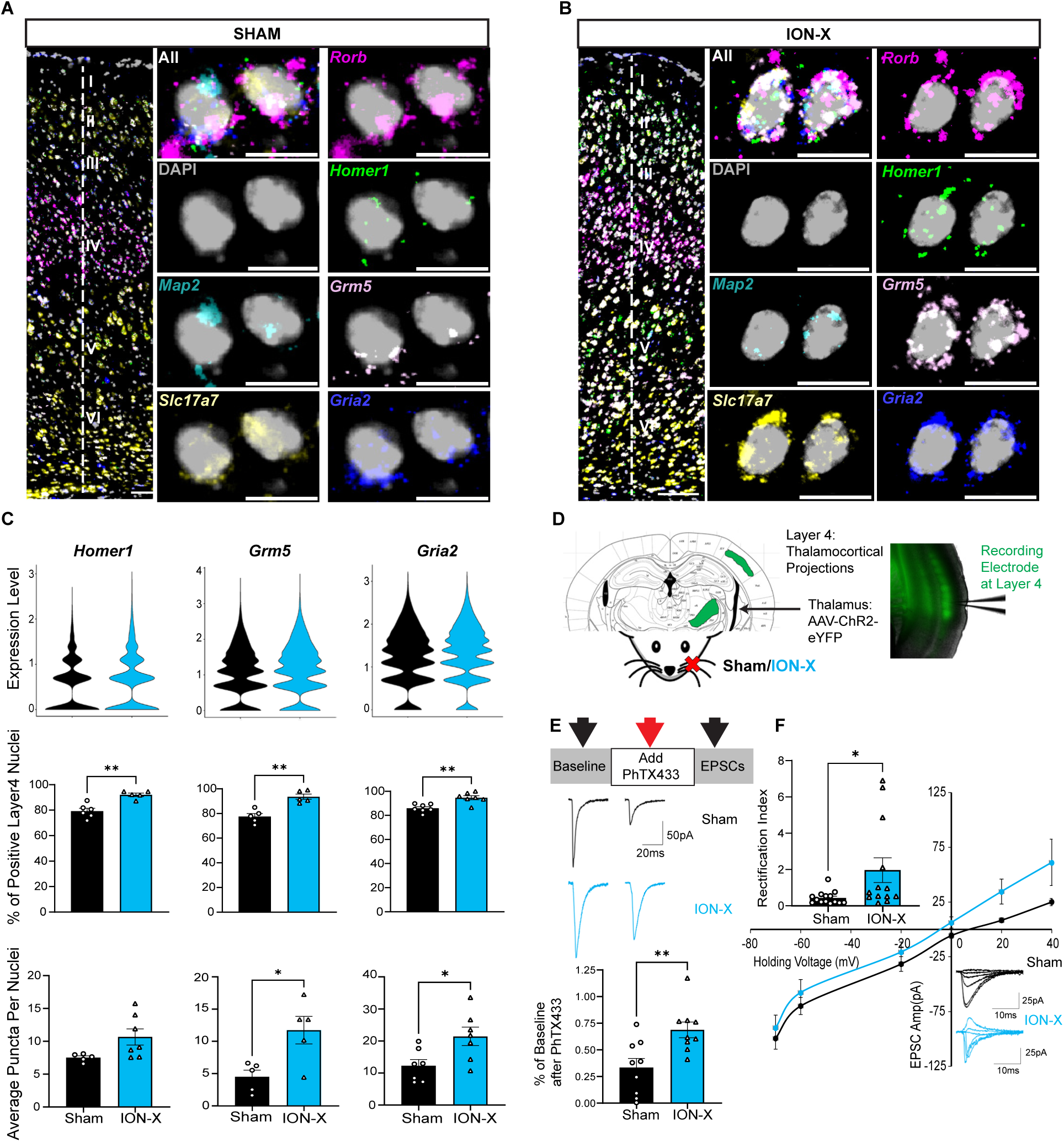
Altered Glutamate Receptor Signaling in the Intact Sensory Pathway after ION-X. A,B) Representative images of mFISH for genes involved in glutamate receptor signaling: *Homer1, Grm5, and Gria2* in sham (left) and ION-X (right) in L4 excitatory neurons expressing *Map2, Slc17a7 and Rorb*. Scale bars for cortical layers (left) 100µm and nuclei (right) at 10µm. C) Violin plots (top) of snRNA-seq predicted changes in gene expression after ION-X and quantification of mFISH images (bottom two). D) AAV-Channel Rhodopsin (ChR2) with an eYFP reporter enabled quantification of TC synapses. E) TC synapses were less sensitive to PhTX433 after ION-X. F) TC synapses had an increased rectification index in L4 neurons after ION-X. See also Table S7, S8.

Given that most mRNAs undergo multiple modifications before being translated to synaptic proteins, we tested if the upregulated gene expression for Gria2 yielded more GluA2-containing AMPARs in L4 excitatory neurons through whole-cell electrophysiology. An adenoassociated virus expressing Channelrhodopsin with an eYFP reporter (AAV9-Syn-ChR2-eYFP) was injected into the sensory thalamus (ventroposteromedial nucleus, VPM) and L4 excitatory neurons were patched for recording (**Figure 3D**). After establishing a stable baseline to excitatory post synaptic current (EPSC) response to LED stimulation of ChR2-expressing TC synapses, a selective antagonist of GluA2-lacking AMPARs (Philanthotoxin, PhTX433) was applied to the bath. After 20 minutes of PhTX433 wash-on, the EPSC amplitude in ION-X showed less sensitivity to PhTX433 compared to shams (**Figure 3E,Table S8**) (t-test p = 0.006). The decreased sensitivity of TC synapses to PhTX433 in ION-X indicates that TC to L4 connections contain more GluA2-containing AMPARs after ION-X ^24,25^. The presence of GluA2-containing AMPARs confers calcium impermeability and can be detected by an increase in the rectification index: the ratio of EPSC amplitude evoked at positive versus negative holding potentials^26^. TC synapses to L4 excitatory neurons had a higher rectification index in ION-X compared to shams (**Figure 3F,Table S8**) (t-test p = 0.044). The inward rectification and higher PhTX433 sensitivity in shams is consistent with previous reports of GluA2-lacking AMPARs in S1BC under control conditions^27^. Together, these results indicate that the increased expression of Gria2 mRNA yields an increased presence of functional GluA2-containing AMPARs at TC synapses after ION-X. GluA2-containing AMPARs are believed to stabilize recently potentiated synapses^28^. Our findings demonstrate that whisker denervation induces significant transcriptional and functional changes in L4 excitatory neurons receiving TC inputs, highlighting the importance of glutamatergic transmission in compensatory plasticity changes within the intact cortex after unilateral whisker loss.

### Increased Synaptogenesis in Intact Layer 4 Neurons After ION-X

Following unilateral ION-X, L4 excitatory neurons responding to the intact whisker set display significantly upregulated gene expression related to synaptogenesis. We focused on four genes that were upregulated in snRNA-seq: *Epha7, Ptn, Pcdh15, Cdh12* (adjusted p = 1.3×10^-4^, 2.9×10^-17^, 1.4×10^-18^, 0.02, respectively) (**Figure 2D, 4**). Ephrins and their receptors support the developmental maturation of synaptic connections^29^*. Epha7* in particular plays a critical role in synaptic refinement as neurons establish functional synapses^30^. Pleiotrophin (*Ptn*) regulates neural outgrowth and is known to modulate plasticity^31,32^. *Pcdh15* and *Cdh12* encode cadherin proteins that promote calcium-dependent cell-cell adhesion^33^, which may also enhance activity-dependent synaptic formation. mFISH was performed in intact S1BC and quantifying the gene expression patterns in *Map2/Slc17a7/Rorb* positive nuclei detected increased expression in ION-X compared to shams (**Figure 4A-D, Table S7; Figure 4B**, *Epha7;* % positive nuclei: t-test p=0.0143; *Ptn*; % positive nuclei: t-test p=0.0086; average puncta p=0.0069; **Figure 4D**, *Pcdh15;* average puncta: t-test p=0.0112; *Cdh12*; % positive nuclei: t-test p=0.0144). These data indicate that snRNA-seq detected increases in gene expression related to synaptogenesis can also be confirmed by mFISH.

**Figure 4.**
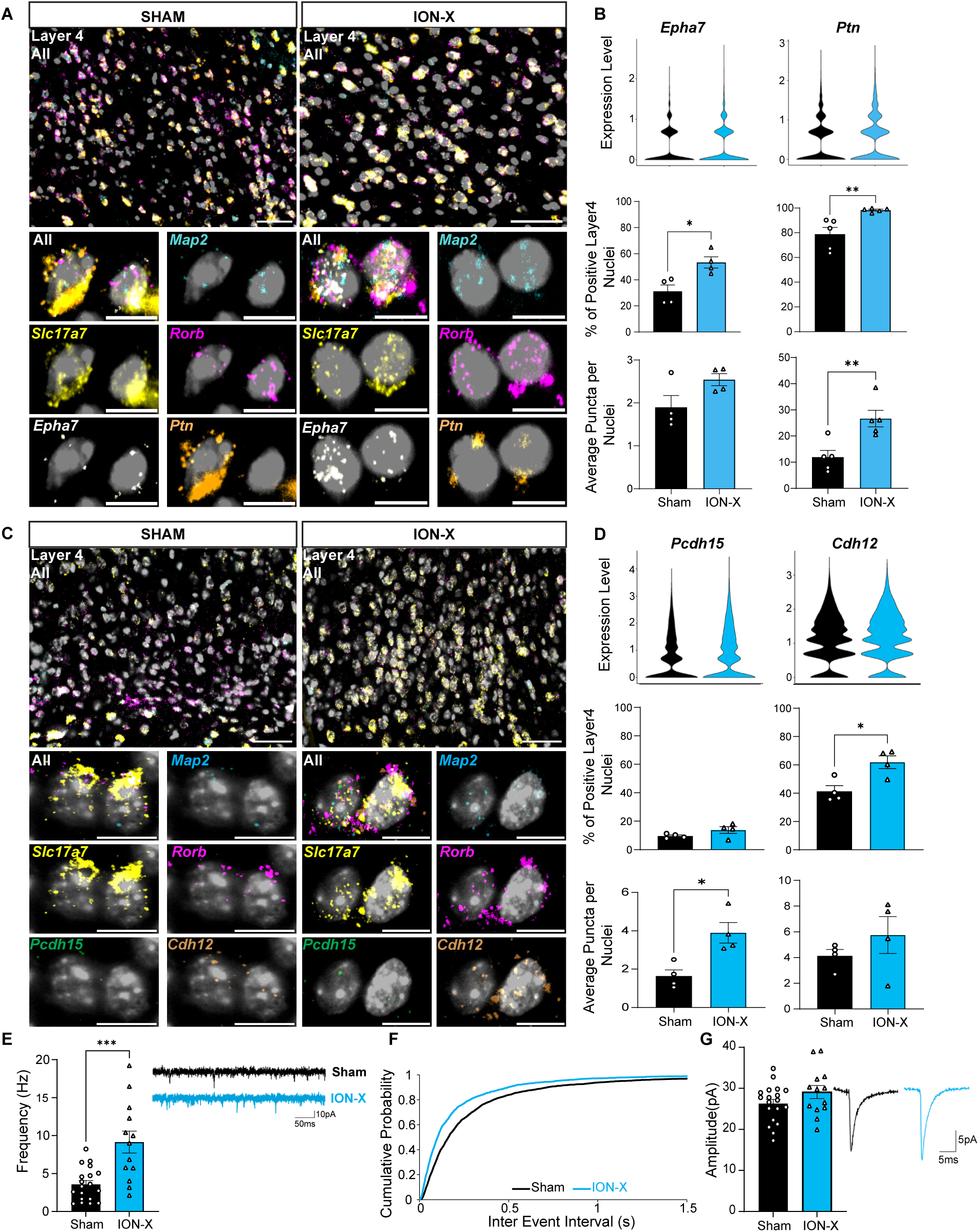
Increased Synaptogenesis in Intact Layer 4 Neurons After ION-X. A) Representative images of mFISH for synaptogenesis genes: *Epha7* and *Ptn* in sham (left) and ION-X (right) samples. Scale bars at 50µm, top and lower at 10µm. B) Violin plots of snRNA-seq predicted changes in gene expression after ION-X and quantification of mFISH images for *Epha7* and *Ptn*. C) Representative images of mFISH synaptogenesis mRNA: *Pcdh15* and *Cdh12* in sham (left) and ION-X (right) samples. Scale bars at 50µm for top and lower at 10µm. D) Violin plots of snRNA-seq predicted changes in gene expression after ION-X and quantification of mFISH for *Pcdh15* and *Cdh12.* E) Spontaneous excitatory post-synaptic currents (sEPSCs) had increased frequency, F) with shorter inter-event-intervals G), but no change in amplitude after ION-X. See also Tables S7,S8.

Next, to evaluate if the increase in synaptogenesis genes yielded more functional synaptic connections, L4 excitatory neurons were targeted for whole-cell recordings and spontaneous EPSCs (sEPSCs) were measured. After ION-X, sEPSC frequency was significantly increased (t-test p = 0.003), (**Figure 4E-F**,**Table S8**), with no change in amplitude (t-test p = 0.15) (**Figure 4G, Table S8**). Previous findings in this model detected increased TC evoked strontium-mEPSC amplitudes after ION-X^19^, however we did not detect increased sEPSC amplitude. That these additional inputs may arise from connections including, but not limited to, the known potentiated TC synapse^17^. The increased sEPSC frequency demonstrates that there are more functional excitatory connections to L4 excitatory neurons after ION-X. Although an increase in frequency could also suggest changes in the release probability of presynaptic partners^34^, the elevated synaptogenesis gene expression point to an increase in the number of synapses as a likely dominant mechanism. The upregulation of these genes, all key players in synapse formation suggests increased excitatory activity and connectivity within L4 neurons. The changes in gene expression and synaptic adaptations likely contribute to the reorganization of TC and cortical circuits, facilitating functional plasticity in the intact pathway after unilateral sensory loss.

## Discussion

There has been a renewed interest in adult TC plasticity after injury that causes a loss in sensory experience^18,35,36^. One hallmark of this plasticity is the potentiation of the intact TC pathway; this likely supports enhanced processing of spared sensory inputs(11, 17). These adaptations include a re-emergence of the molecular mechanisms usually reserved for juvenile circuitry within the critical period^19^. In the spared TC connection, long term potentiation (LTP) occurs in visual system pathways after deafening^35^, and in auditory circuits after vision loss ^17^. Visual TC LTP can also be induced in adult animals with environmental enrichment^37^ or by modulating visual experience^18^, which is mediated by the activity of extracellular matrix-regulating molecules^38^. After unilateral ION-X in adult rats this TC plasticity includes an increase in silent synapses, synaptic number,and TC LTP(19). These changes largely mimic the mechanisms that occur during the critical period of the whisker sensory system^39^. We hypothesized that these changes occur in conjunction with other adaptations that had yet to be discovered. We identified additional plasticity mechanisms with snRNA-seq, mFISH and electrophysiology. This strategy enabled unbiased detection of how TC recipient neurons adapt and influence circuit remodeling after sensory loss. We discovered that after unilateral ION-X, L4 neurons responding to the intact whiskers had differentially expressed genes related to altered glutamate receptor function and synaptogenesis. These transcriptional and functional changes likely support the underlying rejuvenated plasticity to adapt after sensory loss.

### Isolating Layer 4 Excitatory Neurons

Excitatory neurons in L4 of sensory cortices differ based on morphology, electrophysiological properties, and gene expression profiles^40–42^. *Rorb* is a transcription factor that is required for the formation of barrels in L4 of rodent S1BC^23^. When we combined expression of *Rorb* with that of genes for excitatory neurons (*Slc17a7, Map2/Rbfox3*), we successfully isolated L4 excitatory neurons in both snRNA-seq and mFISH experiments. Additional inspection of *Rorb* positive nuclei suggested that additional subcategories of L4 excitatory neurons existed in this data set (**Figure S4**), however further work and a more complicated gene expression configuration may be required to fully characterize these groups. These results highlight the importance of validating snRNA-seq findings with additional complementary techniques^43–45^.

### Glutamate Receptor Signaling

Sensory loss changes how the brain processes signals from the intact sensory inputs, and these changes are mediated by synaptic plasticity. Genes encoding for glutamate receptors and their interacting proteins were altered in L4 excitatory neurons, including *Gria2* (GluA2)*, Grm5* (mGluR5), and *Homer1*. Dynamic control of AMPARs enable synapses to regulate synaptic strength ^46^. The presence of the GluA2 subunit in AMPARs renders these ionotropic glutamate receptors calcium impermeable, and this type of AMPAR swaps in to replace GluA2-lacking AMPARs to stabilize recently potentiated synapses^28,46^. Here an increase in *Gria2* mRNA and GluA2-containing AMPARs after ION-X suggests that the intact TC pathway likely stabilizes recently potentiated feed-forward signals from the intact whiskers; this likely compensates for the loss of contralateral whisker sensation. This conclusion aligns with previous findings that LTP occurs at this synapse 2 weeks after ION-X^19^ which yields yield increased BOLD fMRI activity observed in both rodents^11,12^ and humans^20,47^ after unilateral whisker denervation/injury-induced sensory loss. These adaptations have an early LTP component, where silent synapses containing increased GluN2B-containing NMDARs and TC potentiation occur^19^, followed by a late LTP consolidation component, supported by increased prevalence of GluA2-containing AMPARs. SnRNAseq detected additional gene expression changes related to a potentially third phase: homeostatic adaptations^48,49^.

In addition to altered ionotropic AMPAR signaling, we also detected increased expression for *Grm5* (encoding for mGluR5 receptor) and *Homer1* (encoding for scaffolding protein Homer1) in L4 excitatory neurons after ION-X. mGluR5’s perisynaptic location^50,51^ detects glutamate spillover during periods of high activity and regulates downstream plasticity mechanisms^52^. mGluR5s are anchored near the post-synaptic density by Homer 1b/c^53^, and this anchoring is disrupted by Homer1a during periods of high neuronal activity^54^. Although snRNA-seq and mFISH do not discriminate between the downstream Homer1 protein isoforms, the increased *Homer1* expression is likely to be dominated by the immediate early gene *Homer1a*, which would be upregulated in response to increased intact S1BC activity after ION-X^11,19^. The potential roles for Homer1a after ION-X could include neuroprotection against excitotoxicity^55^ or binding to mGluR5 to enhance NMDA receptor function to support LTP^56,57^. Alternatively, Homer1a’s binding to mGluR5 causes constitutive activation of mGluR5 while simultaneously inhibiting NMDAR function^58,59^. Such inhibition could provide negative feedback to homeostatically reduce global (within the neuron) synaptic strength in response to the increased spontaneous excitatory transmission and the potentiated TC connection^19^ from the intact whiskers^49,60^. Thus, an increase in mRNA levels for *Homer1* and *Gmr5* may stabilize LTP along the intact TC pathway following unilateral whisker denervation in addition to preserving neuronal activity within a physiologically normal range.

### Synaptogenesis

L4 excitatory neurons had upregulated synaptogenesis pathways, and subsequent mFISH detected increased mRNA for cell-cell synaptic adhesion including *Epha7, Pcdh15,* and *Cdh12*^61^. Additionally, *Ptn* (encoding for pleiotrophin), promotes neurite outgrowth in juvenile neural circuits^31^. These pathways trigger increased numbers of synaptic connections between L4 excitatory neurons and other cells after ION-X, and was confirmed with a higher spontaneous mEPSC frequency. Synaptogenesis has important implications for recovery after injury. For example, after unilateral motor cortex stroke, synaptogenesis in neurons within healthy but stroke-adjacent regions and contralateral cortex correlates with enhanced recovery of the affected limb^62,63^. Unilateral sensory loss recruits new innervation from neighboring cortical regions ^64–67^ which may occur in our model of unilateral whisker denervation. Indeed, this type of synaptic rewiring is thought to enhance recovery by making new connections to overcome stroke^68^ or other injuries^69,70^. Connectivity between L4 barrels in S1BC is rare, so the increased synaptogenesis likely arrives either through a post-critical period recruitment of TC inputs, or via other inputs in the cortical column. In support of the increased TC synaptogenesis, this connection has an increased prevalence of GluN2B-containing NMDA receptors, which are components of the silent synapses that precede mature adult synapses^19^. The synaptogenesis observed here reflects initially juvenile, but newly formed synapses that support enhanced processing of the intact whisker set following the loss of the contralateral set.

### Additional Unexplored Avenues for Plasticity

Experience-dependent plasticity triggers altered gene expression to support circuit refinement^71–73^. We only explored a few of the 403 genes that were upregulated in L4 excitatory neurons after ION-X. Other genes of interest include those related to multiple molecular pathways important for plasticity. For example, Neuregulin 3 (*Nrg3*) communicates with ErbB4 tyrosine kinase receptors to regulate synapse formation^74^ which inhibits ocular dominance plasticity^75^. In our dataset, the increased *Nrg3* expression may support the consolidation of TC LTP, thus locking in the stronger intact TC pathway to enhance activity driven by the intact whisker set. Additionally, histone deacetylase (HDAC) activity is thought to be important in regulating memory-enabling long-term plasticity^76^, for example histone H3 phosphorylation and H3/H4 acetylation are high in young animals and stimulation of those pathways can increase ocular dominance plasticity^77^. After ION-X, L4 excitatory neurons have significantly increased gene expression of *Hdac9*, which regulates dendrite development and is upregulated during periods of high activity^78,79^. Here, increased *Hdac9* gene expression may support synaptogenesis and the formation of new connections after unilateral whisker loss. Finally, Ca^2+^/calmodulin-dependent protein kinases (CAMK) comprises a family of molecules that are triggered by activity and signal for synaptic LTP^80,81^. In L4 excitatory neurons after ION-X, gene expression of *Camk2d, Camk1,* and *Camk4* are all significantly increased, indicating that potentiation of synaptic inputs onto these neurons trigger downstream signaling cascades to support adaptations of whisker loss.

### Implications for Recovery after Sensory Loss

Strengthening responses to intact senses after the loss of others may support recovery after injury. In particular, strengthening the feed-forward TC synapses would amplify the sensory signal into the cortex, enhancing the perception of intact incoming sensations^17^. After unilateral whisker denervation, the intact whisker set becomes crucial for rodents to navigate their environment. This may be analogous to individuals learning to rely on their intact limbs to perform daily tasks after unilateral limb loss^1^. In this study, we have determined additional plasticity mechanisms that likely increase activity in the sensory cortex circuitry after unilateral sensory loss. It is unknown if this increased activity enhances sensory perception^5,21^, or impairs motor recovery^47,82,83^. Our findings reveal critical molecular and synaptic changes that enhance cortical plasticity in adults, providing a basis to explore strategies to harness these plasticity mechanisms to support compensatory plasticity and adaptive circuit remodeling after injury.

## Resource availability

Lead Contact:

Requests for further information and resources should be directed and will be fulfilled by Emily Petrus.

## Materials availability

This study did not generate new unique reagents. The mice and reagents used in this study are all commercially available as described in the methods section.

## Data and code availability

Data generated by single nucleus RNA sequencing is available via the NCBI-based Gene Expression Omnibus and will be available upon publication and request. R-based code for analyses will be available in a publicly available repository that is linked to this publication’s DOI. Any additional information required to reanalyze the data reported in this paper is available from the lead contact upon request. mFISH and electrophysiology data will be shared by the lead contact upon request.

## Acknowledgments

The authors wish to thank Kathy Sharer and Abdel Elkalouhn for single nucleus RNA sequencing sample preparation and processing. We thank Ariel Levine and Hey-Kyoung Lee for helpful discussions and comments. We also thank Dennis McDaniel from the USUHS Bioinstrumentation Core for microscopy support. Finally, we thank the USUHS Student Bioinformatics Initiative and Alex DeCasien for bioinformatics support. K.J and A.P.K were supported by the intramural research program of the NINDS, NIH.

## Author Contributions

L.I. and E.P performed wet lab experiments, data analysis, and wrote the manuscript. S.C. and K.J. performed data analysis. E.P and A.P.K. provided scientific and technical guidance throughout the project and feedback on the written manuscript.

## Supplemental Information

Document S1: Figures S1-S4 and Table S7-S9.

Tables S1-S6: Excel files containing data too large to fit in a PDF, related to Figures 1, 2.

## STAR METHODS

### Key Resources Table

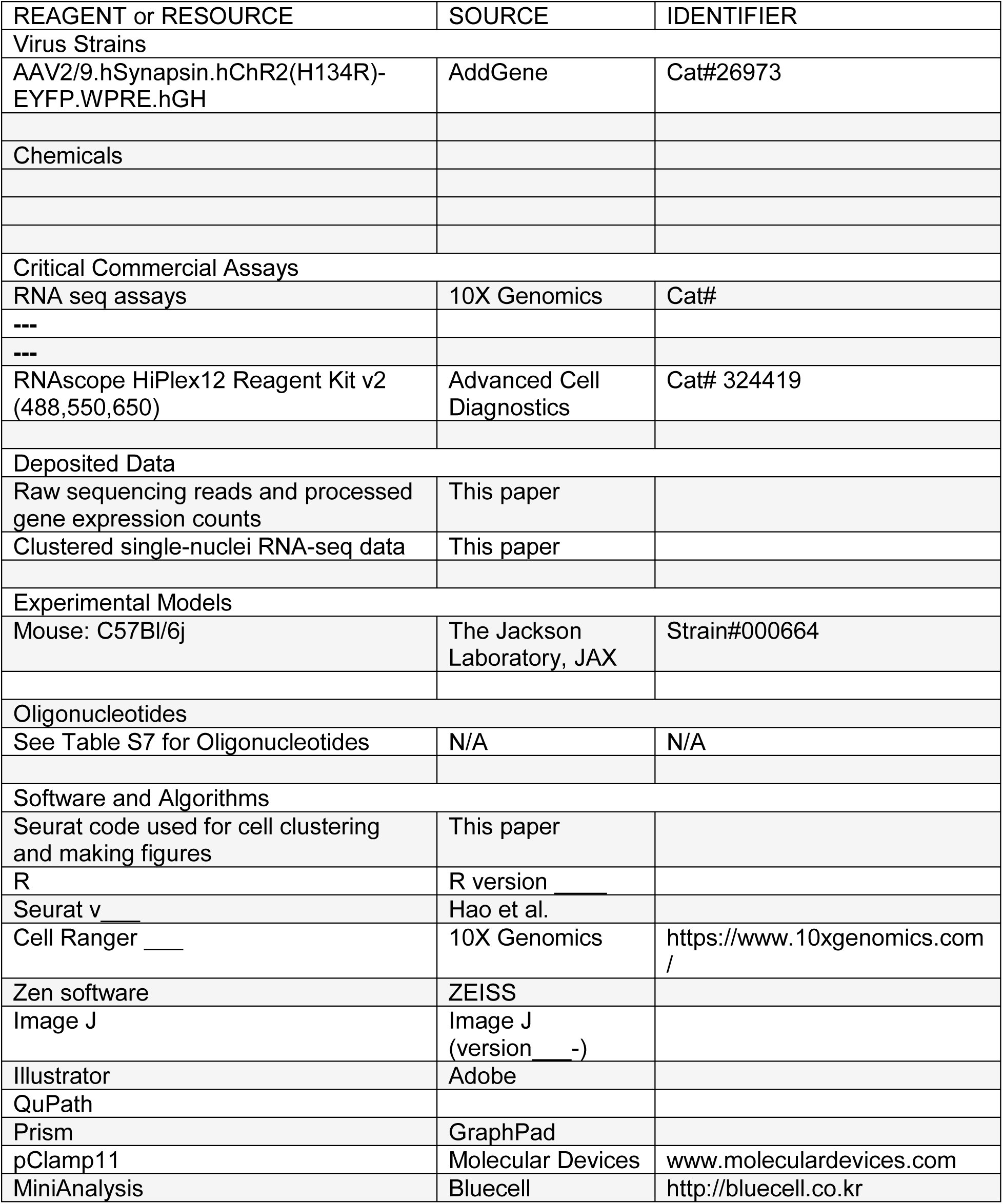

| REAGENT or RESOURCE | SOURCE | IDENTIFIER |
| --- | --- | --- |
| Virus Strains |  |  |
| AAV2/9.hSynapsin.hChR2(H134R)-EYFP.WPRE.hGH | AddGene | Cat#26973 |
| Chemicals |  |  |
| Critical Commercial Assays |  |  |
| RNA seq assays | 10X Genomics | Cat# |
| --- |  |  |
| --- |  |  |
| RNAscope HiPlex12 Reagent Kit v2 (488,550,650) | Advanced Cell Diagnostics | Cat# 324419 |
| Deposited Data |  |  |
| Raw sequencing reads and processed gene expression counts | This paper |  |
| Clustered single-nuclei RNA-seq data | This paper |  |
| Experimental Models |  |  |
| Mouse: C57Bl/6j | The Jackson Laboratory, JAX | Strain#000664 |
| Oligonucleotides |  |  |
| See Table S7 for Oligonucleotides | N/A | N/A |
| Software and Algorithms |  |  |
| Seurat code used for cell clustering and making figures | This paper |  |
| R | R version ____ |  |
| Seurat v ____ | Hao et al. |  |
| Cell Ranger ____ | 10X Genomics | https://www.10xgenomics.com / |
| Zen software | ZEISS |  |
| Image J | Image J (version ____ -) |  |
| Illustrator | Adobe |  |
| QuPath |  |  |
| Prism | GraphPad |  |
| pClamp11 | Molecular Devices | www.moleculardevices.com |
| MiniAnalysis | Bluecell | http://bluecell.co.kr |

## EXPERIMENTAL MODEL AND SUBJECT DETAILS

### Animals

All procedures were approved by the National Institutes of Health Animal Care and Use committee (ACUC) under protocol 1160, and the Uniformed Services of the Health Sciences Institutional Animal Care and Use Committee (IACUC) under protocol APG-22-081. Facilities in both universities are accredited by the association for assessment and accreditation of laboratory animal care (AAALAC). C57Bl/6j mice (Strain#000664, RRID: <u>IMSR_JAX:000664</u>) were obtained from Jackson Laboratories (Bar Harbor, ME) and arrived on site at 3-5 weeks of age. SnRNA-seq experiments included only male mice but subsequent multiplex fluorescent *in situ* hybridization (mFISH) and electrophysiological experiments were male/female balanced. No statistically significant differences were identified between genders. Littermates were housed 2-4 per cage with food and water ad libitum on a 12-hour light/dark cycle.

### Surgery

All surgeries were performed under aseptic techniques. After each surgery, animals received Meloxicam slow release to combat inflammation. Animals remained on a heated surface and returned to the animal facility after they were fully ambulatory, typically within 20-60 minutes.

#### Stereotaxic Surgery

At 3-4 weeks of age, mice were anesthetized with 1-3% isoflurane mixed with O_2_ and unilaterally injected (left side) with 500nl of a virus encoding for channel rhodopsin (ChR2): AAV2/9.hSynapsin.hChR2(H134R)-EYFP.WPRE.hGH (RRID: <u>Addgene_26973</u>, titre 2.1×10^13^). Animals received the viral injection into the sensory thalamus: ventral posteromedial nucleus (VPM). Coordinates were ML 1.6, AP 2.0, Depth 3.5. The virus was delivered with a 2.5µl Hamilton syringe with a 30G beveled needle. Injection duration was 10 minutes with the needle left in place for 10 additional minutes to ensure adequate perfusion into the tissue. The incision was closed with tissue staples. These mice were used for electrophysiology recordings.

#### Whisker Denervation Surgery (ION-X)

After one week of habituation to in-house animal facilities, 4-6-week-old mice were anesthetized with an intraperitoneal (IP) injection of a ketamine/xylazine/saline cocktail (80mg/kg ketamine, 10mg/kg xylazine). Once mice were unresponsive to paw pinch, the right side of the face was shaved and an incision caudal to the whisker pad was made to visualize the infraorbital nerve (ION) bundle. For sham mice, the procedure was concluded at this step, with the incision closed with a single suture and a drop of tissue glue (Tissuemend II, Veterinary Product Laboratories). In ION-X mice, the ION bundle was cut with scissors and the incision was closed as described above. Animals then received an IP injection of Antisedan (1mg/kg; Atipanezole, Zoetis) to reverse the anesthesia.

### Single-nucleus RNA sequencing

The protocol was followed according to Sathyamurthy et al 2018^84^.

#### Sample Collection and Processing

12 days after sham/ION-X surgery, male mice were anesthetized with 5% isoflurane until the absence of a righting reflex was observed. Mice were then decapitated and the intact (contralateral to the intact whiskers) S1BC was rapidly dissected into ice-cold dissection buffer (80 mm NaCl, 3.5 mm KCl, 1.25 mm H2PO_4_, 25 mm NaHCO_3_, 4.5 mm MgSO_4_, 0.5 mm CaCl_2_, 10 mm glucose, and 90 mm sucrose), which was bubbled continuously with a 95% O_2_/5% CO_2_ gas mixture. Two S1BCs (one per mouse) were collected per sample, with an average collection time of 2-4 minutes. Each sample was denounced using five strokes with Pestle A, followed by five strokes with pestle B (Kimble: Kontes Dounce Tissue Grinder) in sucrose buffer containing the following: 0.32 M sucrose, 10mM HEPES (pH 8.0) 5mM CaCl_2_, 3mM Mg-acetate, 0.1mM EDTA, 1mM Dithiothreitol, 0.1% Triton X-100. The lysate was diluted with 3mL of sucrose buffer and centrifuged at 3,200xg for 10 minutes. The supernatant was removed and 3mL sucrose buffer was added, followed by a 2 minute incubation. The pellet and sucrose were transferred to an Oak Ridge centrifuge tube, where the pellet was homogenized using an Ultra-Turrax (IKA Instruments, Germany) for 1 minute at setting 1. Then 12 mL of density buffer (1M sucrose, 10mM HEPES, 3mM Mg-acetate, 1mM Dithiothreitol) was added below the layer of nuclei. The tube was centrifuged again at 3,200xg for 20 minutes. The supernatant was poured off, and the nuclei were collected from the walls of the tube with 1mL PBS plus 0.02% Bovine Serum Albumin (BSA). This solution was spun once more at 3,200xg for 10 minutes. Nuclei were resuspended in PBS with 0.02% BSA. Samples were placed on ice and transferred to the National Human Genome Research Institute single cell genomics core facility for processing on 10X and Illumina platforms. 8 samples (4 sham, 4 ION-X, 2 mice per sample) were sequenced on 3 separate sample dates. The kits used are as follows: 10x Chromium Next Gen Chip G Single Cell Kit, 16rxns; Chromium Next Gem Automated Single Cell 3’ Library & Gel Bead Kit v3.1, 4rxns; Illumina Next-Seq 500/550 High Output Kit v2.5 (150 cycles). High output run with 400M reads.

The 10X Cell Ranger workflow (v7.0.0) was used to align sequences to the reference mouse genome. Single nuclei sequencing is preferable over single-cell for this experiment as we are looking for transcriptional synaptic changes and this technique has less tissue handling and dissociation-related effects leading to a more comprehensive picture of cell type proportions.

#### snRNA-seq data analysis

R (version 4.4.1) was used in conjunction with functions supported in Seurat Hao et al 2021 (version 5.1.0) unless otherwise noted. Filtered “h5” files output from cellranger (Sample Collection and sequencing) were imported into R by sample using the “Read10X_h5” function. The “CreateSeuratObject” function was then applied to these files to produce one Seurat object per sample containing the total number of counts per cell (nCount_RNA) and the total number of detected genes per cell (nFeature_RNA). The “PercentageFeatureSet” function was then applied to each of these objects (pattern = “^mt-”) to enumerate the percent total counts per cell associated with mitochondrial genes (percent.mt). After, these objects were collapsed into one using the “merge” function. The “VlnPlot” function was then applied to this collapsed object to visually compare the percent.mt, nCount_RNA, and nFeature_RNA values across samples (Figure S1). Cells observed to not have a percent.mt < 5, a nCount_RNA > 675, a nCount_RNA < 15000, a nFeature_RNA > 500, and a nFeature_RNA < 5000 were filter removed from further analysis. Post filtering, the collapsed object was split back into individual objects by sample using the “SplitObject” function. Count data contained within each split-out object was then normalized using the “SCTransform” function (vars.to.regress = “percent.mt”, method=”glmGamPoi”). Integration of this now normalized data across objects into a single object was accomplished using the “SelectIntegrationFeatures” function (nfeatures = 3000) and the “PrepSCTIntegration” function followed by use of the “FindIntegrationAnchors” and “IntegrateData” functions (normalization.method = “SCT”). Post integration, the “RunPCA” and “ElbowPlot” functions were applied to generate and explore the percent total variation explained for each component up to 200 components (npcs = 200). From this inspection, 100 components were selected to be used for clustering. For clustering, the “RunUMAP” function was used in conjunction with the “FindNeighbors” function (dims = 1:100) followed by use of the “FindClusters” function over a range of resolutions (0.4:1.5). Clustering results were then inspected at each resolution using the “DimPlot” function (group.by=”UMAP”) as well as across resolutions using the “clustree” function. From this inspection, 0.80 was selected as the optimal resolution. To identify and remove doublet cells and doublet clusters at this resolution, the “computeDoubletDensity” (https://plger.github.io/scDblFinder/articles/computeDoubletDensity.html) and “findDoubletClusters” (https://plger.github.io/scDblFinder/articles/findDoubletClusters.html) functions were used. A z-score was also calculated for each cell by cluster using its UMAP1 value in conjunction with the mean (trim=0.20) and SD calculated from all UMAP1 values for cells that fall in the same cluster. Cells observed to have a z-score > 2 were then filter removed as cluster-level outliers. This same procedure was repeated using UMAP2 values. Surviving cells per cluster were then summarized by sample via stacked bar plot using the “ggplot” and “geom_bar” functions (position=”stack”). For cluster annotation, the “RunAzimuth” function was used (reference = “mousecortexref”). To test for and identify markers for each cluster, the “FindConservedMarkers” function was used. Test results were then inspected and putative marker genes selected to augment the Azimuth assigned cell type. To visualize marker expression, both the “FeaturePlot” and “VlnPlot” (stack=TRUE) functions were used. To rank prioritize clusters on differences occurring between sample conditions (i.e., ION-X vs Sham), the “augur” function (https://github.com/neurorestore/Augur) was used. The above steps post integration were repeated two separate times on two cluster subsets: 1) those observed to have positive expression of Rbfox3, Map2, or Slc17a7 (i.e., excitatory neurons) and 2) those assigned as being type L4 neuron. For 1), the number of components used was 50 with 0.04 selected as the optimal resolution. For 2), the number of components used was 100 with 0.80 selected as the optimal resolution.

Differentially expressed genes from L4 neurons with an adjusted p-value of 0.05 or less were used to conduct Gene Ontology enrichment analysis using the clusterProfiler (4.62) package for R (3.16.0). We compared our gene list to the mouse gene database -org.Mm.eg.db-to identify overrepresented biological processes, molecular functions, and cellular compartments. The top 10-11 entries for each category are listed by q-value and colored according to the number of DEGs included^85^.

### Multiplex Florescent In situ Hybridization (mFISH)

#### Sample Preparation

12 days after sham/ION-X surgery, C57BL6/J mice were anesthetized using isoflurane and perfused with ice-chilled 1X PBS followed by 4% paraformaldehyde (PFA). Whole brains were removed and immersed in 4%PFA for 24hrs. They were then sequentially incubated for 24hrs in 10%, 20%,and 30% sucrose and embedded in OCT, frozen and stored at −80°C. Frozen S1BC sections of 12µm were cut using cryostat (CM1950 Leica Biosystems) and collected onto SuperFrost Plus Slides. To prevent RNA degradation, the slides were immediately frozen at - 20°C. To validate the results from the snRNA-seq experiments and to visualize multiple gene candidates in one slice, we performed HiPlex12 Reagent RNAScope Assay (Advanced Cell Diagnostics,Cat. 324419) according to the manufacturer’s protocol. The list of oligonucleotides used for target probes is in the Table S9. The sections were briefly rinsed with PBS, baked at 60°C, re-perfused with PFA, and dehydrated through graded ethanol concentrations. The slides were then immersed in a target retrieval buffer in a steamer for 5 minutes and dried overnight circled with a hydrophobic barrier. The next day, slides were incubated with Protease III reagent and then subsequently with a pooled set of 12 HiPlex probes for 2 hours at 40°C. Probes with different tails attached (T1-T12) allowed for specific and accurate detection of the desired gene. Sections were then treated with three amplifying solutions (Amp 1-3) for 30 minutes each, followed by 3% FFPE reagent to reduce autofluorescence. The last step was the addition of fluorophore for the corresponding channels (488, 550, 650nm) which allowed for the detection of 3 targeted mRNA molecules in each round. DAPI was used as a counterstain and tiled images were captured on Zeiss Axioscan 3.0 microscope with a 40X objective.

After each round, the sections were cleaved using a 10% cleaving solution, and the next set of detection reagents was applied. At the end of 4th round, the sections were cleaved again and a blank image was captured without any probes for background subtraction. All the sets of images from each round for each section were aligned and registered to HiPlex Image Registration Software v2.1.0 (ACDBio) and a final composite image with 12 probes was generated. Each probe was stained in 5-6 mice.

### Electrophysiological Recordings

#### Acute slice preparation

Two weeks after ION-X or sham surgeries acute slices were made for whole-cell electrophysiological recordings. Mice were anesthetized using isoflurane vapors (5% mixed with O_2_) until the absence of the corneal reflex was observed. The brain was quickly dissected and immersed in ice-cold dissection buffer (80 mm NaCl, 3.5 mm KCl, 1.25 mm H_2_PO_4_, 25 mm NaHCO_3_, 4.5 mm MgSO_4_, 0.5 mm CaCl_2_, 10 mm glucose, and 90 mm sucrose), which was bubbled continuously with a 95% O_2_/5% CO_2_ gas mixture. Brain blocks containing primary somatosensory cortex were dissected and coronally sectioned into 300-µm-thick slices using a Leica VT1000S vibratome (Leica Biosystems Inc.). Slices were incubated for 60 minutes at room temperature before recordings began.

#### Recordings

Slices were transferred to a submersion-style recording chamber mounted on a fixed stage (Sutter Instruments Company) with an upright Nikon Eclipse FN1 microscope (Nikon Instruments) and illuminated with oblique infrared (IR) illumination. Recordings were performed in artificial CSF (ACSF; 130 mm NaCl, 3.5 mm KCl, 1.25 mm NaH_2_PO_4_·H_2_O, 24 mm NaHCO_3_, 10 mm glucose, 2.5 mm CaCl_2_, and 1.5 mm MgCl_2_) bubbled with 95% O_2_/5% CO_2_ at 30°C.

0.005mM bicuculline and 0.1 mM APV were added to ACSF to isolate AMPA receptor-mediated excitatory events. The ACSF was perfused at a rate of 2 ml/min. Voltage clamp experiments used Cs-gluconate internal solution, containing the following: 130 mm Cs-gluconate, 8 mm KCl, 1 mm EGTA, 10 mm HEPES, 4 mm ATP, 5 mm QX-314; pH 7.3, 285–295 mOsm. Neurons in L4 were targeted for patch clamp recordings. with an access resistance higher than 25 MΩ and input resistance lower than 100 MΩ were discarded. Cells which had more than a 20% fluctuation in these values for recordings were excluded from analysis. A maximum of three cells per animal, one cell per slice were recorded for each dataset.

#### *Rectification Index* 0.1mM spermine (Tocris) was added to the internal solution

Monosynaptic EPSCs were evoked by 455nm LED illumination of ChR2-expressing TC terminals. Light intensity was adjusted so events were approximately 100pA at −70mV. EPSCs were recorded in pseudorandom order at holding potentials including: −70, −60, −20, 0, +20 and +40mV at 0.1Hz with 6-10 events per holding potential. The rectification index was calculated as EPSC amplitudes at +40mV/-60mV. 6 Sham and 6 ION-X mice were used to obtain 12 and 13 cells respectively.

#### Philanthotoxin Sensitivity

Monosynaptic EPSCs were evoked by 455nm LED illumination of ChR2-expressing TC terminals. Light intensity was adjusted so events were about 100pA. A 5-minute baseline at 0.1Hz was recorded before the addition of 10µM philanthotoxin 433 (PhTX433) (Aobius, Gloucester MA). EPSCs were recorded at 0.1Hz for up to 30 minutes after PhTX addition. The percent reduction was calculated as amplitude of post-PhTX EPSC/pre-PhTX EPSC. 5 Sham and 6 ION-X mice were used to obtain 9 and 9 cells respectively.

#### Spontaneous Excitatory Post Synaptic Currents (sEPSCs)

Neurons in L4 were targeted for voltage clamp recording and sEPSCs were recorded for 3-5 minutes. ACSF contained 20µM bicuculline and 100µM DL-2-amino-5-phosphonopentanoic acid (DL-APV). 200 consecutive sEPSCs were selected for analysis. 7 Sham and 5 ION-X mice were used to obtain 19 and 13 cells respectively.

## QUANTIFICATION AND STATISTICAL ANALYSIS

SnRNAseq data was analyzed as described in the snRNA-seq data analysis section.

### Analysis of RNA mFISH probes

The signal from the composite image obtained from the ACD HiPlex Registration software was quantified utilizing QuPath (0.4.3) bioimage software. For each section, three non-overlapping annotated regions of S1BC were selected and averaged for the counts. DAPI channel was utilized for nuclei detection, setting the signal threshold above the background intensity for accurate detection of individual nuclei. The subcellular detection module was utilized for probe detection. The dot identification in each channel for individual probes was performed by adjusting their detection threshold value based on their pixel intensity. A nucleus was considered positive or negative for a probe based on the presence of puncta detected^86^. QuPath provided a table with cell-by-cell measurements of all the 12 targeted probes in each nucleus.

The positive expression of *Map2*, *Slc17a7,* and *Rorb* probes were used to identify L4 excitatory neurons, with similar expression levels for these 3 transcripts observed in sham and ION-X samples. Quantification for L4 neurons for other targeted probes was based on the co-expression of the probes with the three marker genes. The probes were quantified in two ways:

1. percentage of nuclei positive for a particular gene: this will determine the total subpopulation of nuclei positive for a particular mRNA transcript out of all the nuclei analyzed (approx. 400 nuclei per annotated region; 1200 per section) in each section 2) average expression of gene per nucleus: this indicates the number of mRNA transcripts in an individual nucleus. This also demonstrates the variability in the expression level of that gene per nuclei. Statistics between the Sham and ION-X samples were performed using t-test and the values are depicted as mean ± SEM.

### Analysis of electrophysiology data

An axon patch-clamp amplifier 700B (Molecular Devices) was used for patch-clamp recordings. Data were acquired through pClamp11 and analyzed with Clampfit 11.2 software (Molecular Devices). sEPSCs were analyzed using MiniAnalysis software by synaptosoft/BlueCell.

Statistics were performed using Student’s t-test between sham vs ION-X. Values are depicted as mean +/- standard error of the mean (SEM).

---

**Figure S1.**
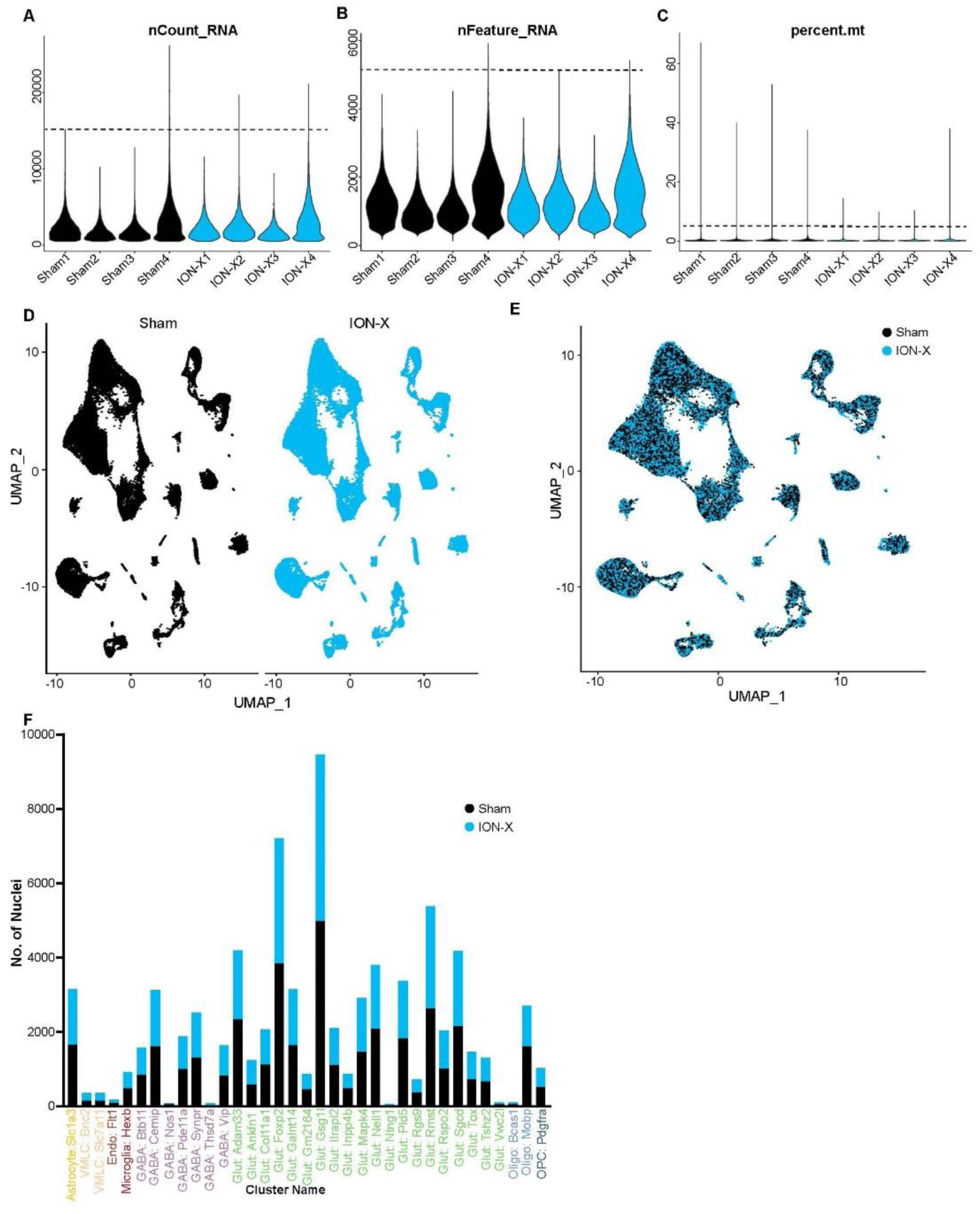
Quality control parameters of the snRNA-seq data set. A-C) Violin plots depicting the number of genes, UMIs, and mitochondrial mRNA detected per sample, with the cut-off marked by the dashed line. D, E) UMAP of the 83,390 nuclei split by the sample. **F**) Bar graph illustrating the distribution of nuclei by sample type.

**Figure S2.**
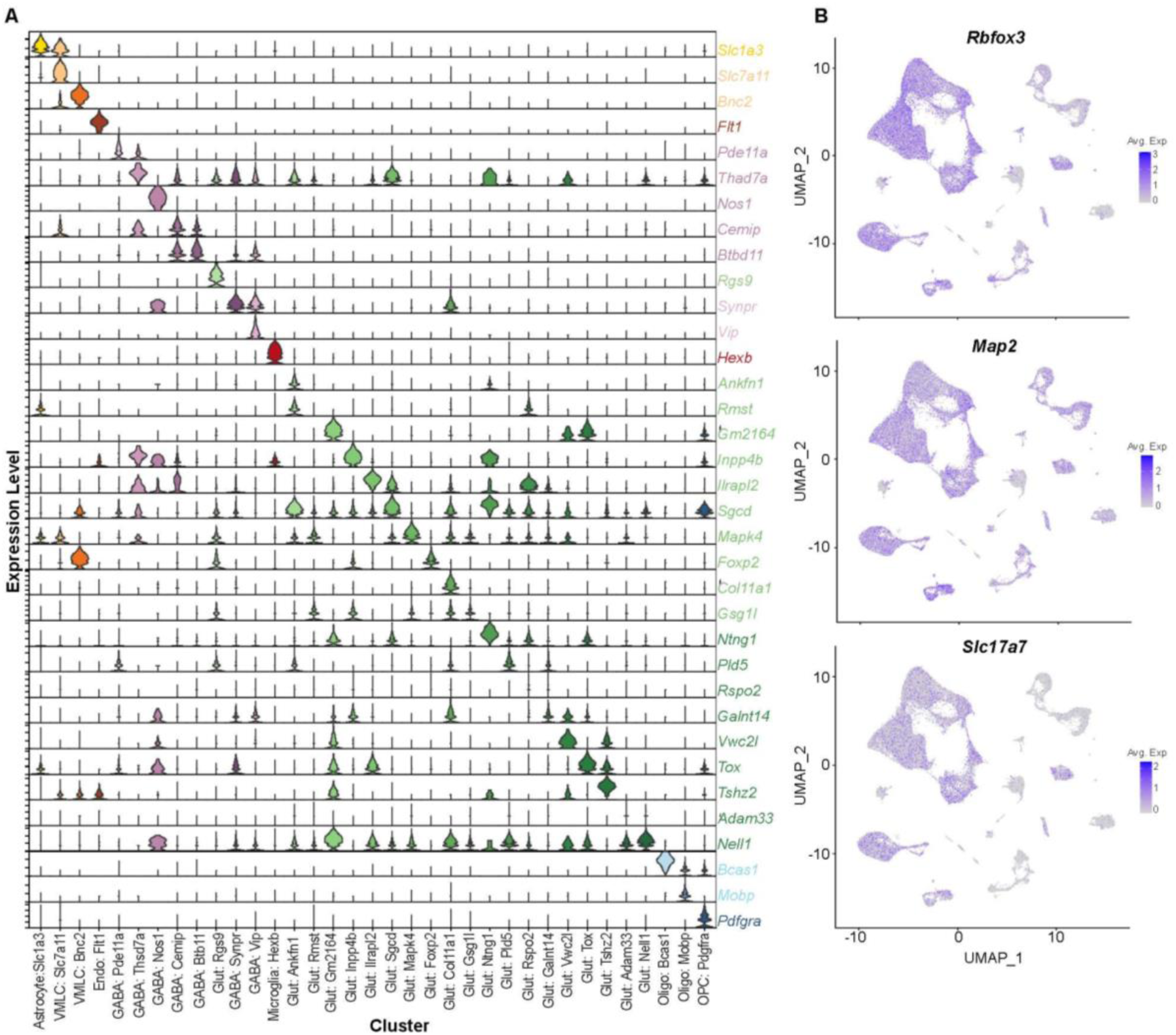
Characterization of the entire S1BC snRNA-seq data set. A) Violin plot of top marker gene for each of the 35 distinct clusters. B) Feature plots of neuronal marker genes Rbfox3, Map2, and excitatory cell marker Slc17a7 in the cell-type based clusters.

**Figure S3.**
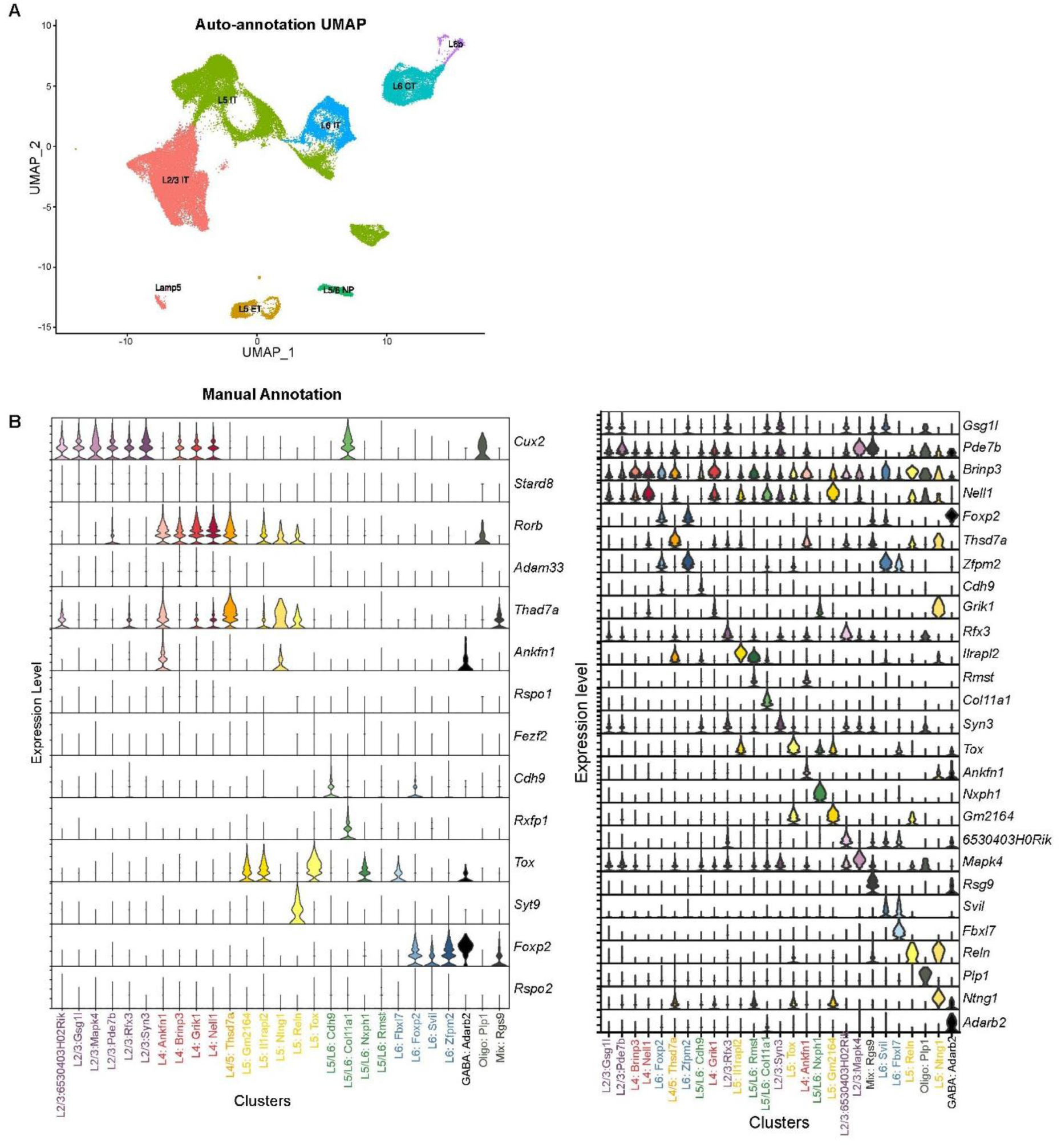
Auto and manual annotation of excitatory neurons. A) Auto annotation UMAP generated with 26 distinct neuron clusters B)Violin plot of manually curated gene markers for different cortical layers (leftand violin plot showing the top gene maker for the manually annoatated 26 clusters found in excitatory neurons from S1BC (right) See also Tables S2,S3.

**Figure S4.**
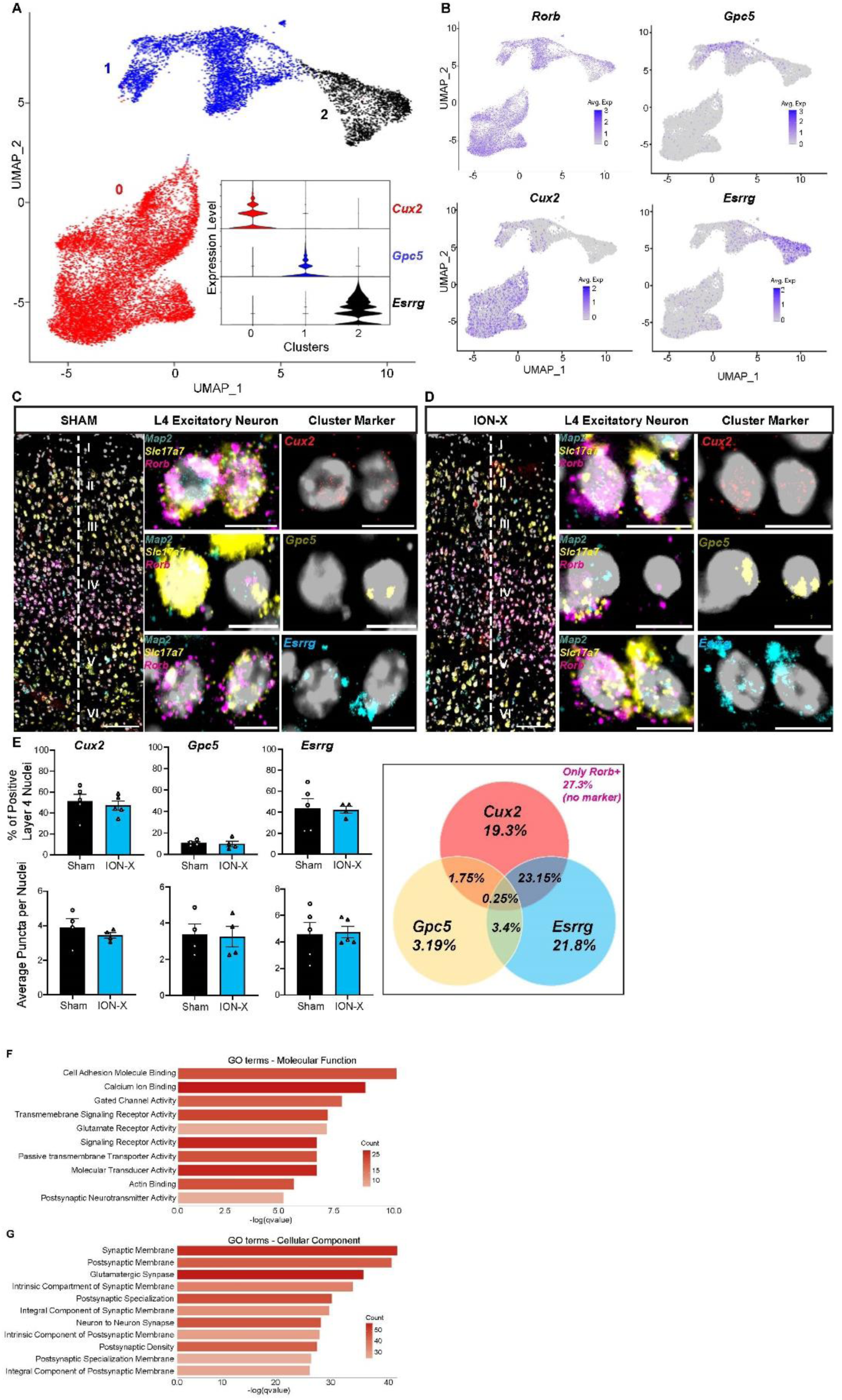
Examination of Layer 4 Excitatory Neurons. A) Re-clustering of L4 neurons reveals 3 distinct clusters, with expression of specific marker genes for each cluster. B) Feature plots with gene expression of each cluster marker (*Cux2, Gpc5, Esrrg*) and *Rorb.* C) Representative sham and ION-X images of mFISH for L4 excitatory neurons (*Map2/Slc17a7/Rorb*) with their cluster markers *Cux2* for Cluster 0, *Gpc5* for Cluster 1, and *Esrrg* for Cluster 2. Scale bars at 100µm for layer images and others at 10µm. E) Gene expression quantification for the cluster makers between sham and ION-X samples and Venn Diagram showing the percentage distribution and overlap between the cluster makers. F, G) Gene Ontology enrichment analysis on collapsed L4 excitatory neuron clusters identified upregulated molecular function and cellular component listed by q-value and colored according to the number of DEGs. See also Tables S4,S5,S6.

**Supplementary Tables.** Supplementary Tables S1-6 are excel files

**Supplementary Table S7.**
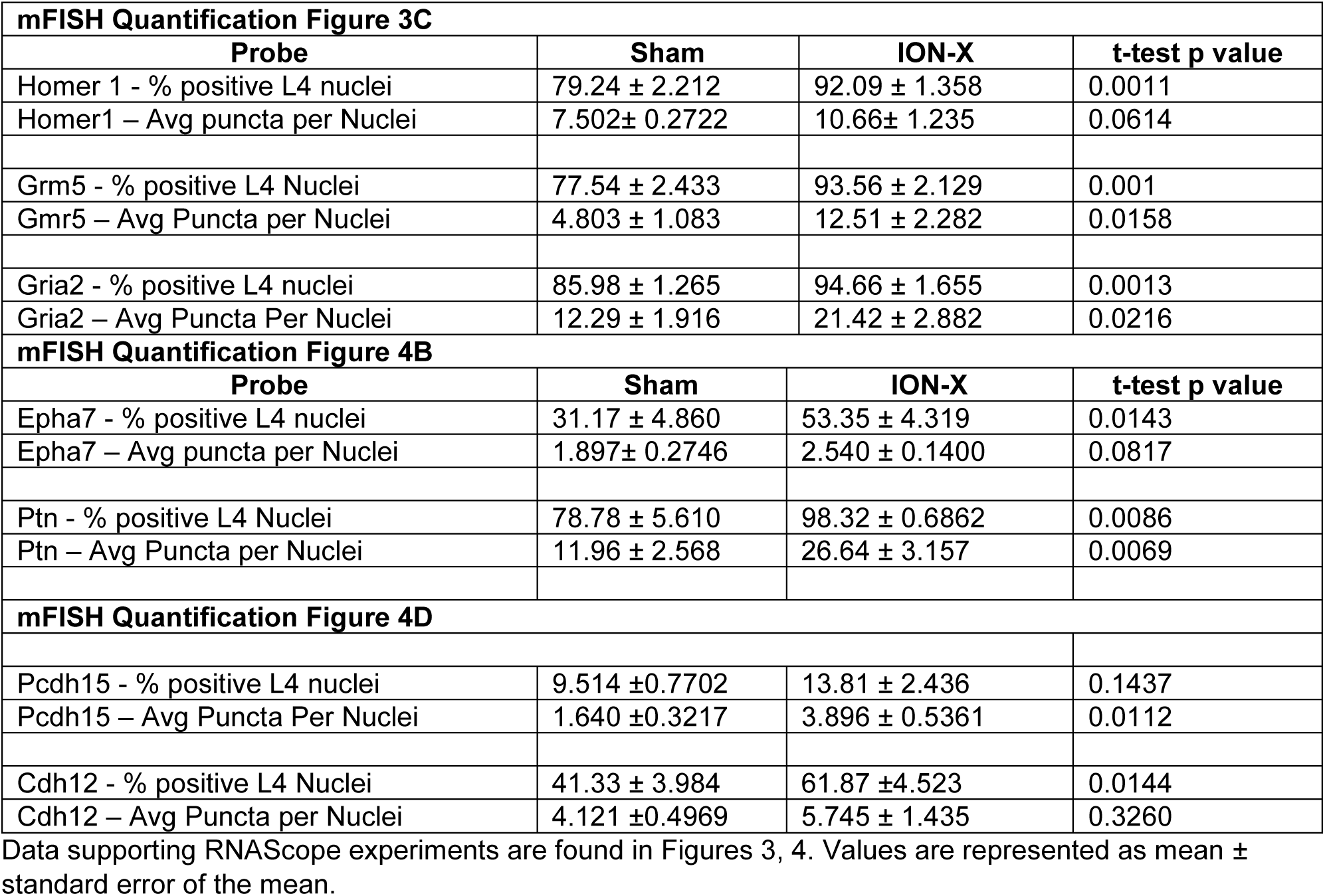
Data supporting RNAScope experiments are found in Figures 3, 4. Values are represented as mean ± standard error of the mean.

**Supplementary Table S8.**
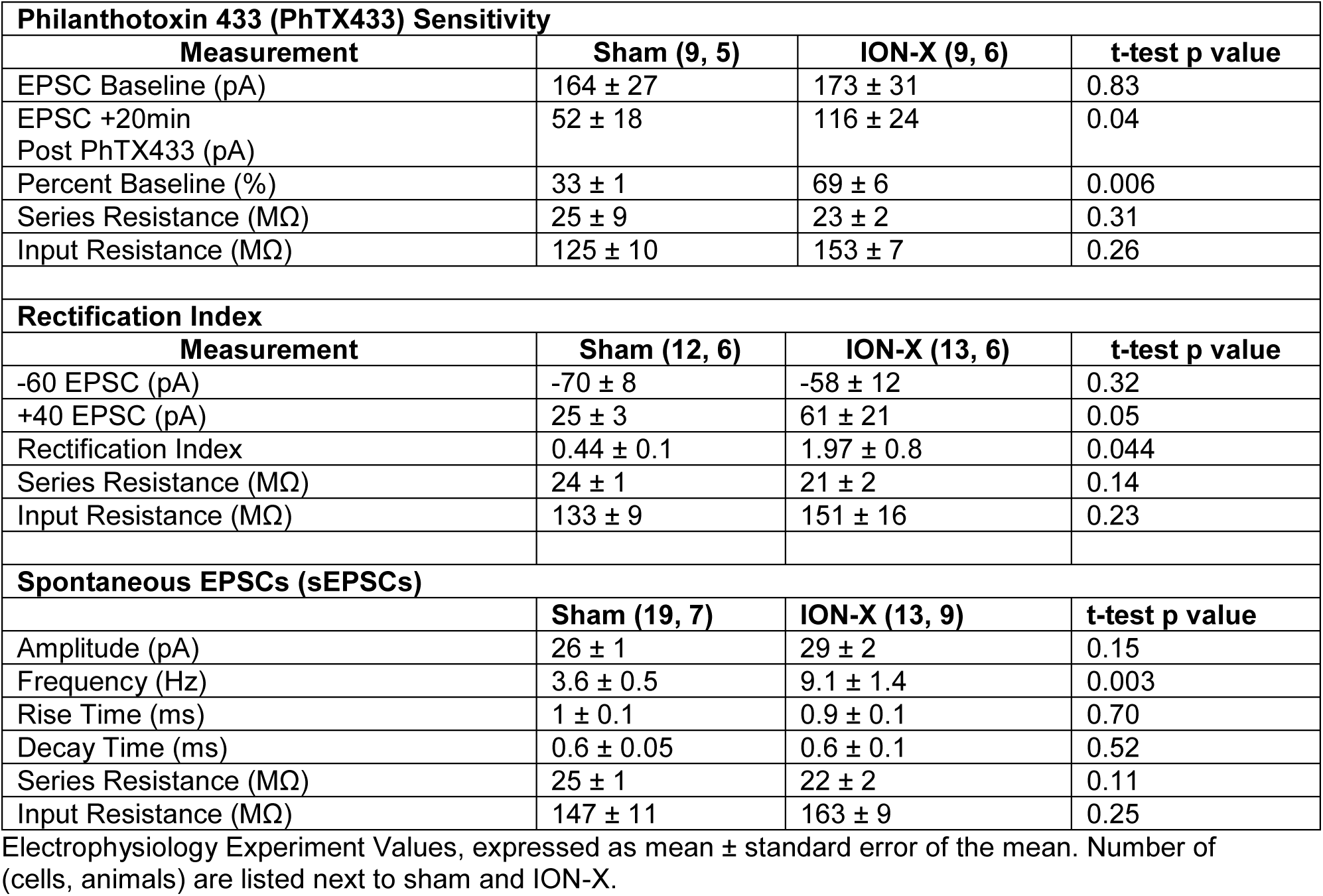

**Supplementary Table S9.**
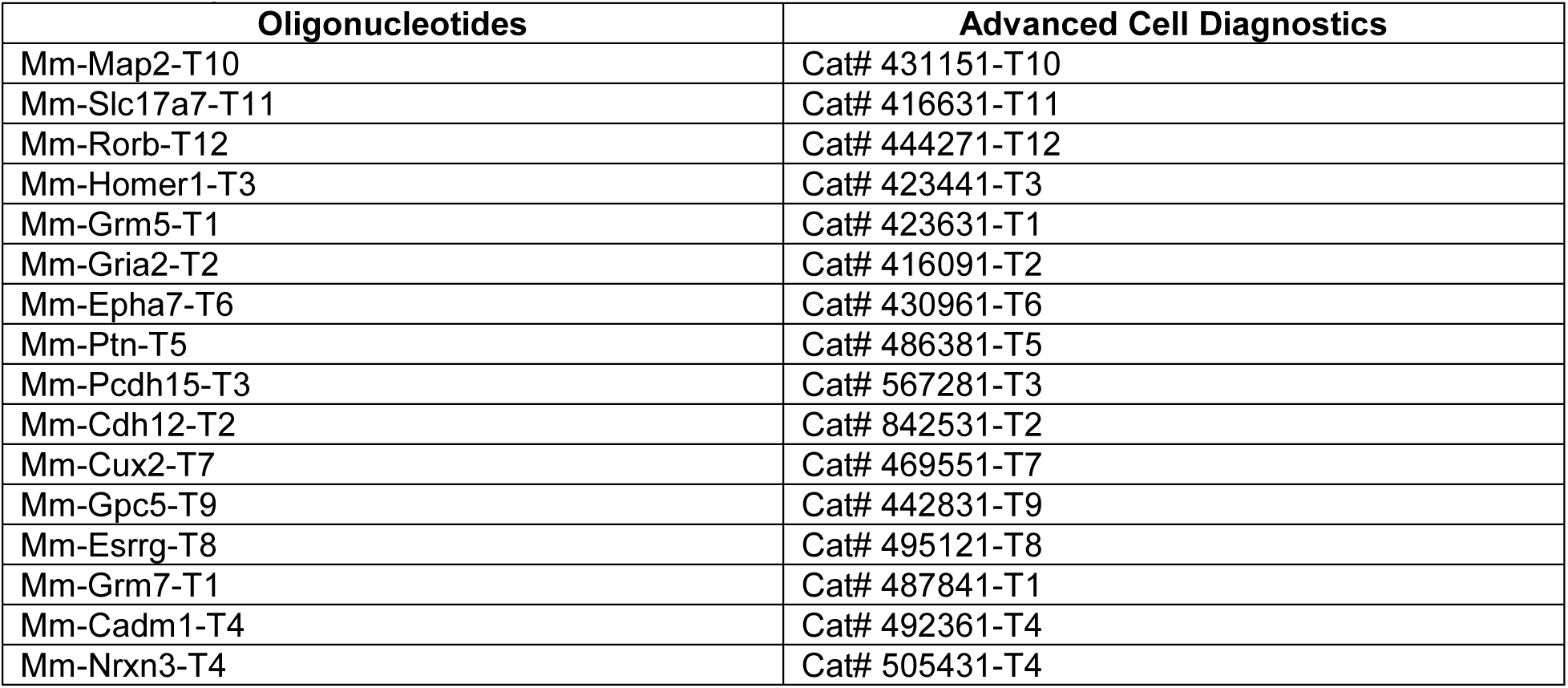

