## Supplemental Figures 1-4, Tables 7-9 for "Post-Critical Period Transcriptional and Physiological Adaptations of Thalamocortical Connections after Sensory Loss"

**Supplemental Information for IyerPetrus_2024**

### Supplementary Figures & Legends

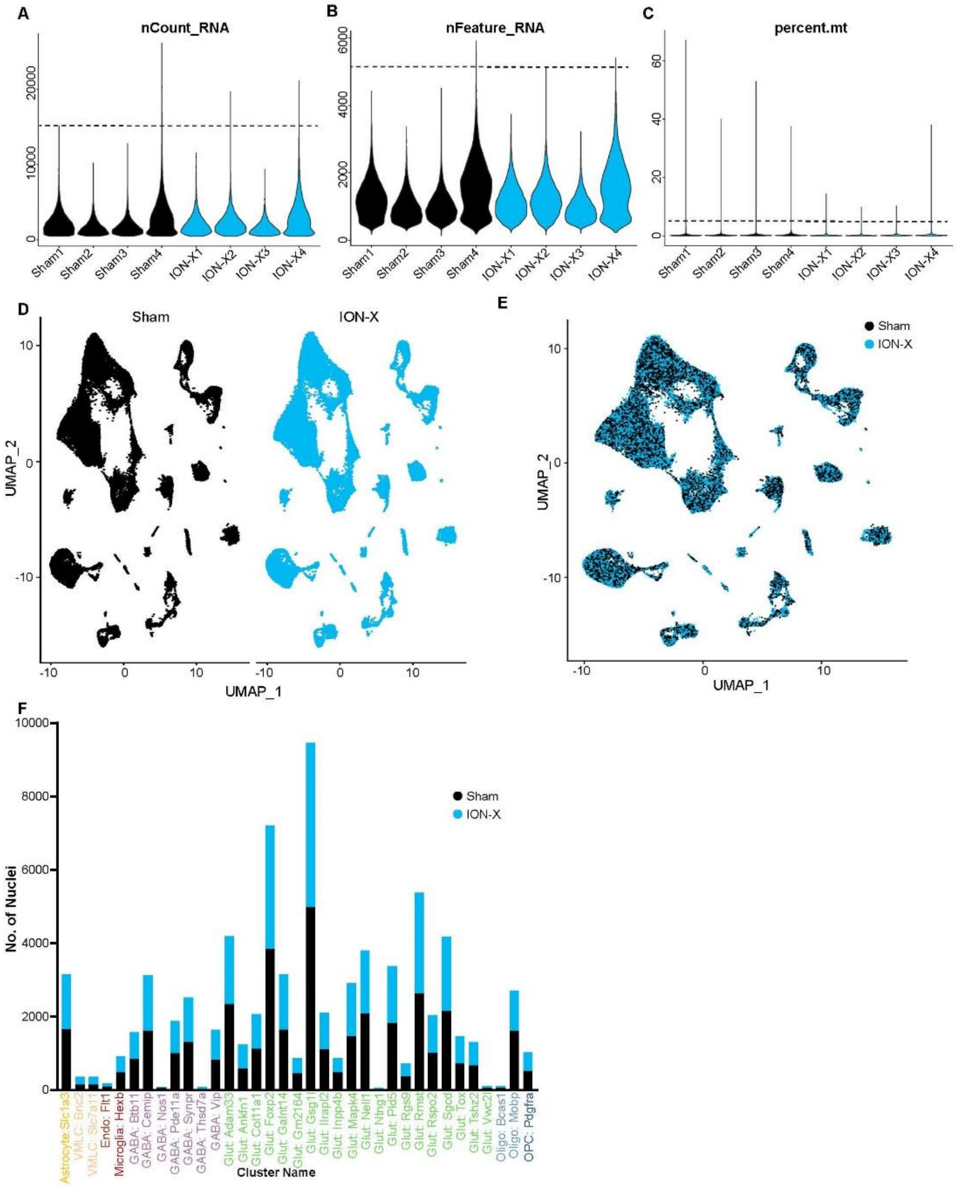

**Figure S1.**
Quality control parameters of the snRNA-seq data set
A-C) Violin plots depicting the number of genes, UMIs, and mitochondrial mRNA detected per sample, with the cut-off marked by the dashed line. D, E) UMAP of the 83,390 nuclei split by the sample. **F**) Bar graph illustrating the distribution of nuclei by sample type.

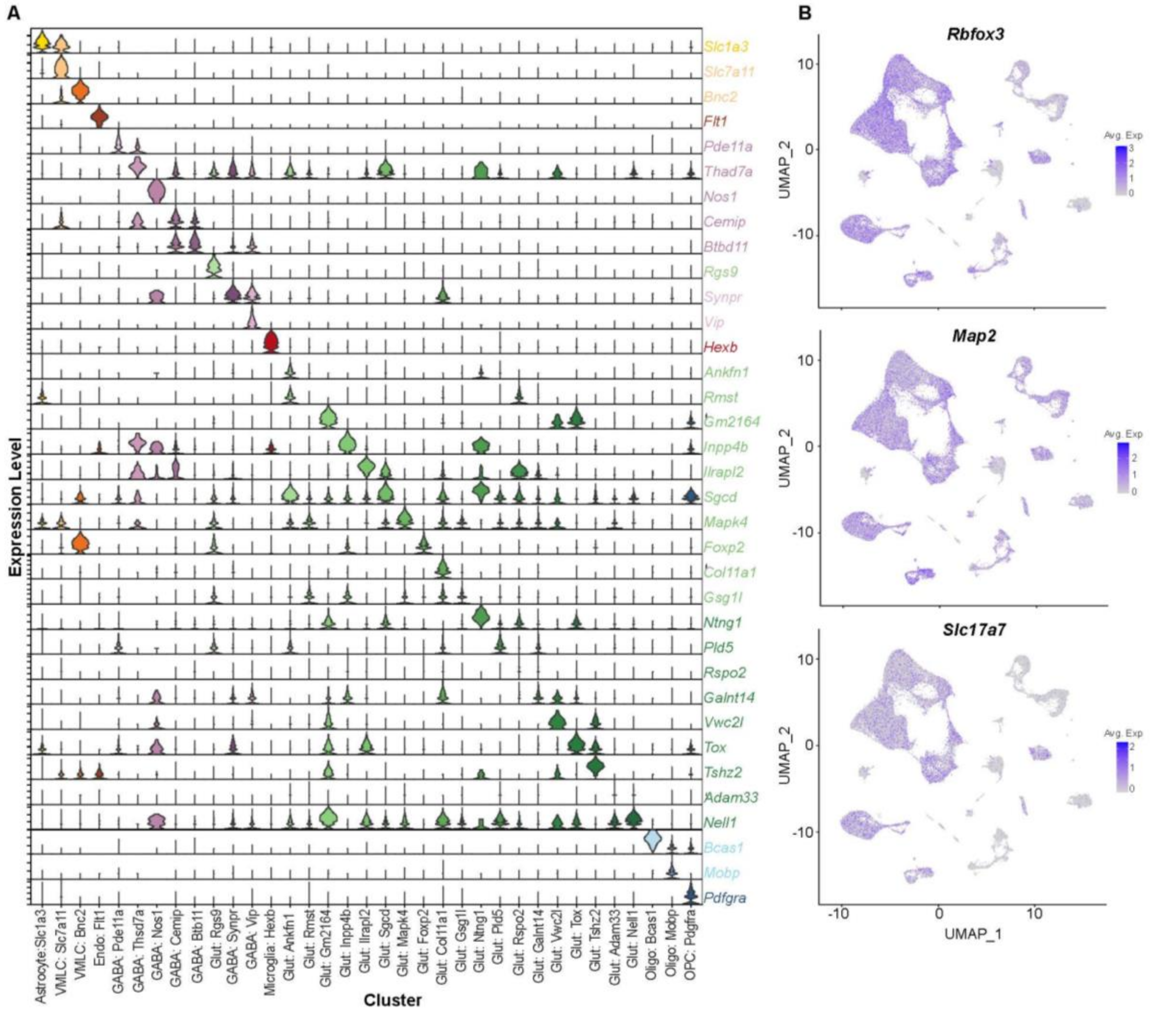

**Figure S2.**

Characterization of the entire S1BC snRNA-seq data set

A) Violin plot of top marker gene for each of the 35 distinct clusters. B) Feature plots of neuronal marker genes Rbfox3, Map2, and excitatory cell marker Slc17a7 in the cell-type based clusters.

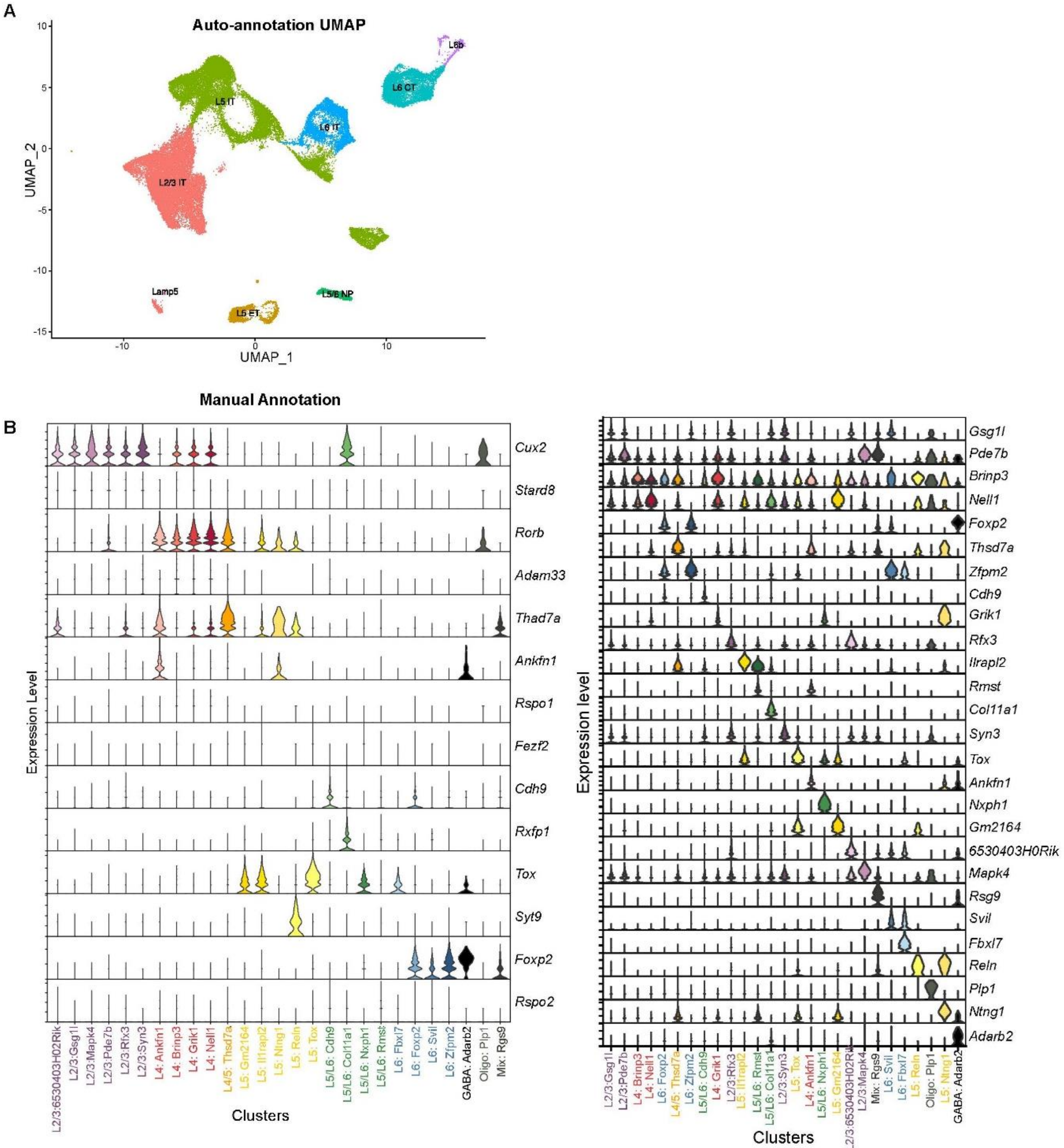

**Figure S3.**

Auto and manual annotation of excitatory neurons

A) Auto annotation UMAP generated with 26 distinct neuron clusters B)Violin plot of manually curated gene markers for different cortical layers (leftand violin plot showing the top gene  maker for the manually annoatated 26 clusters found in excitatory neurons from S1BC (right) See also Tables S2,S3.

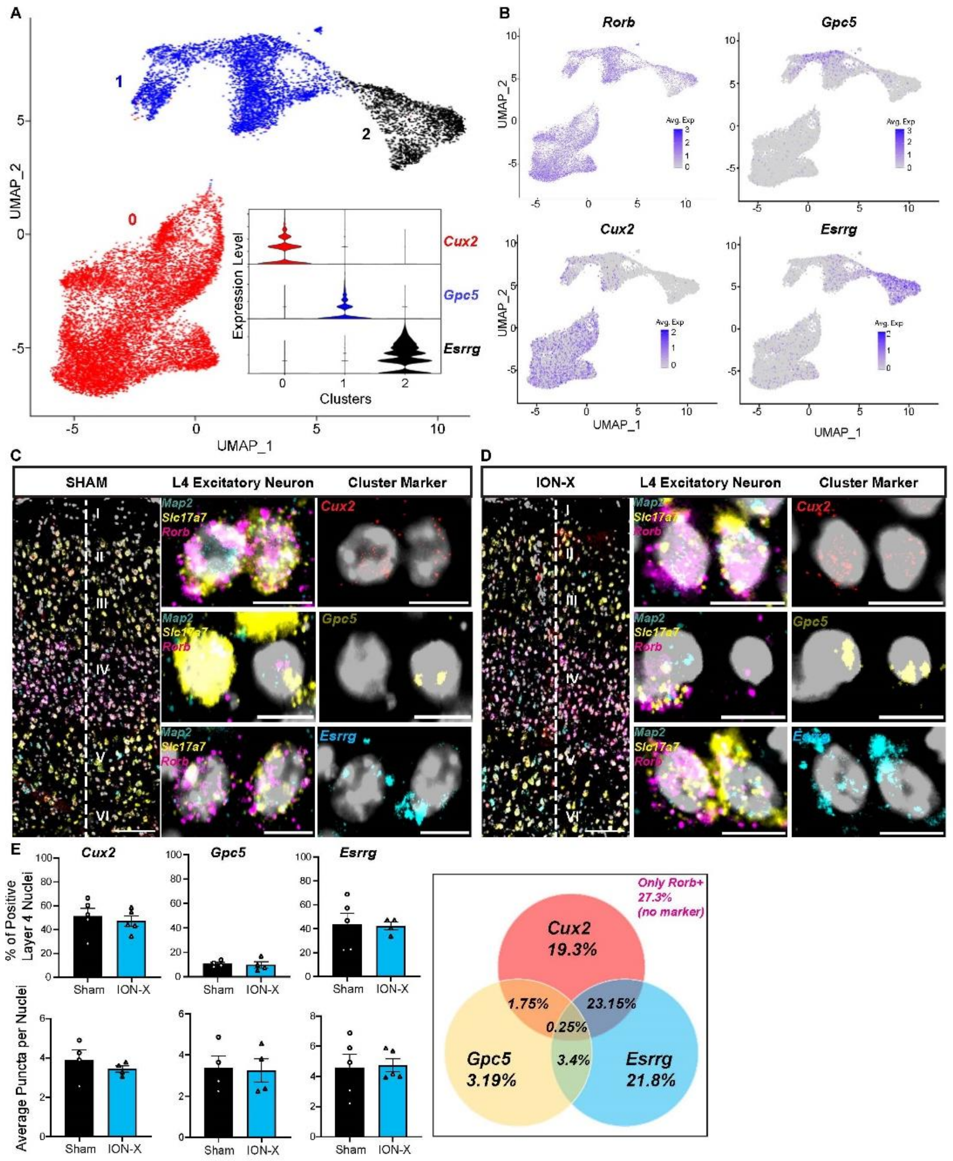

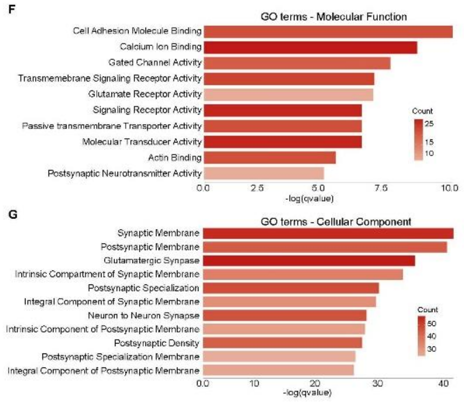

**Figure S4.**

Examination of Layer 4 Excitatory Neurons

A) Re-clustering of L4 neurons reveals 3 distinct clusters, with expression of specific marker genes for each cluster. B) Feature plots with gene expression of each cluster marker (*Cux2, Gpc5, Esrrg*) and *Rorb.* C) Representative sham and ION-X images of mFISH for L4 excitatory neurons (*Map2/Slc17a7/Rorb*) with their cluster markers *Cux2* for Cluster 0, *Gpc5* for Cluster 1, and *Esrrg* for Cluster 2. Scale bars at 100µm for layer images and others at 10µm. E) Gene expression quantification for the cluster makers between sham and ION-X samples and Venn Diagram showing the percentage distribution and overlap between the cluster makers. F, G) Gene Ontology enrichment analysis on collapsed L4 excitatory neuron clusters identified upregulated molecular function and cellular component listed by q-value and colored according to the number of DEGs. See also Tables S4,S5,S6.

**Supplementary Tables**
Supplementary Tables S1-6 are excel files

Supplementary Table S7

| **mFISH Quantification Figure 3C** | | | |
|---|---|---|---|
| **Probe** | **Sham** | **ION-X** | **t-test p value** |
| Homer 1 - % positive L4 nuclei | 79.24 ± 2.212 | 92.09 ± 1.358 | 0.0011 |
| Homer1 – Avg puncta per Nuclei | 7.502± 0.2722 | 10.66± 1.235 | 0.0614 |
| Grm5 - % positive L4 Nuclei | 77.54 ± 2.433 | 93.56 ± 2.129 | 0.001 |
| Gmr5 – Avg Puncta per Nuclei | 4.803 ± 1.083 | 12.51 ± 2.282 | 0.0158 |
| Gria2 - % positive L4 nuclei | 85.98 ± 1.265 | 94.66 ± 1.655 | 0.0013 |
| Gria2 – Avg Puncta Per Nuclei | 12.29 ± 1.916 | 21.42 ± 2.882 | 0.0216 |
| **mFISH Quantification Figure 4B** | | | |
| **Probe** | **Sham** | **ION-X** | **t-test p value** |
| Epha7 - % positive L4 nuclei | 31.17 ± 4.860 | 53.35 ± 4.319 | 0.0143 |
| Epha7 – Avg puncta per Nuclei | 1.897± 0.2746 | 2.540 ± 0.1400 | 0.0817 |
| Ptn - % positive L4 Nuclei | 78.78 ± 5.610 | 98.32 ± 0.6862 | 0.0086 |
| Ptn – Avg Puncta per Nuclei | 11.96 ± 2.568 | 26.64 ± 3.157 | 0.0069 |
| **mFISH Quantification Figure 4D** | | | |
| Pcdh15 - % positive L4 nuclei | 9.514 ±0.7702 | 13.81 ± 2.436 | 0.1437 |
| Pcdh15 – Avg Puncta Per Nuclei | 1.640 ±0.3217 | 3.896 ± 0.5361 | 0.0112 |
| Cdh12 - % positive L4 Nuclei | 41.33 ± 3.984 | 61.87 ±4.523 | 0.0144 |
| Cdh12 – Avg Puncta per Nuclei | 4.121 ±0.4969 | 5.745 ± 1.435 | 0.3260 |

Data supporting RNAScope experiments are found in Figures 3, 4. Values are represented as mean ± standard error of the mean.

Supplementary Table S8

| **Philanthotoxin 433 (PhTX433) Sensitivity** | | | |
|---|---|---|---|
| **Measurement** | **Sham (9, 5)** | **ION-X (9, 6)** | **t-test p value** |
| EPSC Baseline (pA) | 164 ± 27 | 173 ± 31 | 0.83 |
| EPSC +20min Post PhTX433 (pA) | 52 ± 18 | 116 ± 24 | 0.04 |
| Percent Baseline (%) | 33 ± 1 | 69 ± 6 | 0.006 |
| Series Resistance (MΩ) | 25 ± 9 | 23 ± 2 | 0.31 |
| Input Resistance (MΩ) | 125 ± 10 | 153 ± 7 | 0.26 |
| **Rectification Index** | | | |
| **Measurement** | **Sham (12, 6)** | **ION-X (13, 6)** | **t-test p value** |
| -60 EPSC (pA) | -70 ± 8 | -58 ± 12 | 0.32 |
| +40 EPSC (pA) | 25 ± 3 | 61 ± 21 | 0.05 |
| Rectification Index | 0.44 ± 0.1 | 1.97 ± 0.8 | 0.044 |
| Series Resistance (MΩ) | 24 ± 1 | 21 ± 2 | 0.14 |
| Input Resistance (MΩ) | 133 ± 9 | 151 ± 16 | 0.23 |
| **Spontaneous EPSCs (sEPSCs)** | | | |
| | **Sham (19, 7)** | **ION-X (13, 9)** | **t-test p value** |
| Amplitude (pA) | 26 ± 1 | 29 ± 2 | 0.15 |
| Frequency (Hz) | 3.6 ± 0.5 | 9.1 ± 1.4 | 0.003 |
| Rise Time (ms) | 1 ± 0.1 | 0.9 ± 0.1 | 0.70 |
| Decay Time (ms) | 0.6 ± 0.05 | 0.6 ± 0.1 | 0.52 |
| Series Resistance (MΩ) | 25 ± 1 | 22 ± 2 | 0.11 |
| Input Resistance (MΩ) | 147 ± 11 | 163 ± 9 | 0.25 |

Electrophysiology Experiment Values, expressed as mean ± standard error of the mean. Number of (cells, animals) are listed next to sham and ION-X.

Supplementary Table S9

| **Oligonucleotides** | **Advanced Cell Diagnostics** |
|---|---|
| Mm-Map2-T10 | Cat# 431151-T10 |
| Mm-Slc17a7-T11 | Cat# 416631-T11 |
| Mm-Rorb-T12 | Cat# 444271-T12 |
| Mm-Homer1-T3 | Cat# 423441-T3 |
| Mm-Grm5-T1 | Cat# 423631-T1 |
| Mm-Gria2-T2 | Cat# 416091-T2 |
| Mm-Epha7-T6 | Cat# 430961-T6 |
| Mm-Ptn-T5 | Cat# 486381-T5 |
| Mm-Pcdh15-T3 | Cat# 567281-T3 |
| Mm-Cdh12-T2 | Cat# 842531-T2 |
| Mm-Cux2-T7 | Cat# 469551-T7 |
| Mm-Gpc5-T9 | Cat# 442831-T9 |
| Mm-Esrrg-T8 | Cat# 495121-T8 |
| Mm-Grm7-T1 | Cat# 487841-T1 |
| Mm-Cadm1-T4 | Cat# 492361-T4 |
| Mm-Nrxn3-T4 | Cat# 505431-T4 |
